# Integration of cell wall synthesis activation and chromosome segregation during cell division in *Caulobacter*

**DOI:** 10.1101/2022.11.05.515301

**Authors:** Christopher R. Mahone, Xinxing Yang, Joshua W. McCausland, Isaac P. Payne, Jie Xiao, Erin D. Goley

**Affiliations:** Department of Biological Chemistry, Johns Hopkins University School of Medicine, Baltimore, MD, USA; Department of Biophysics and Biophysical Chemistry, Johns Hopkins University School of Medicine, Baltimore, MD, USA; Hefei National Laboratory for Physical Sciences at the Microscale, School of Basic Medical Sciences, Division of Life Sciences and Medicine, University of Science and Technology of China, Hefei, 230026, China

## Abstract

To divide, bacteria must synthesize and remodel their peptidoglycan (PG) cell wall, a protective meshwork that maintains cell shape. FtsZ, a tubulin homolog, dynamically assembles into a midcell band, recruiting division proteins including the PG synthases FtsW and FtsI. FtsWI are activated to synthesize PG and drive constriction at the appropriate time and place, however their activation pathway remains unresolved. In *Caulobacter crescentus*, FtsWI activity requires FzlA, an essential FtsZ-binding protein. Through time-lapse imaging and single-molecule tracking of *C. crescentus* FtsW and FzlA in perturbed genetic backgrounds, we demonstrate that FzlA is a limiting constriction activation factor that converts inactive, fast-moving FtsW to an active, slow-moving state. We find that FzlA interacts with the DNA translocase FtsK, and place FtsK genetically in a pathway with FzlA and FtsWI. Misregulation of the FzlA-FtsK-FtsWI pathway leads to heightened DNA damage and cell death. We propose that FzlA integrates the FtsZ ring, chromosome segregation, and PG synthesis to ensure robust and timely constriction during *Caulobacter* division.

## Introduction

Bacterial cell division is a robust process, evolved over billions of years, that requires tight regulation of multiple essential events to ensure survival. These events include marking the division site, recruiting division proteins, segregating the chromosome, remodeling and synthesizing cell wall, and separating the daughter cells (Dewachter et al., 2018; Mahone and Goley, 2020). The first step in division is the assembly of FtsZ, an essential and conserved tubulin homolog, into a cytokinetic “Z-ring” at the incipient site of division (Mahone and Goley, 2020; Barrows and Goley, 2021). Once the Z-ring is established, dozens of proteins, coined the divisome, are directly or indirectly recruited by FtsZ to the division site (Mahone and Goley, 2020; McQuillen and Xiao, 2020). FtsZ polymers within the Z-ring are highly dynamic, driven by FtsZ’s GTPase activity, and exhibit treadmilling motion about the inner circumference at the division site (Bisson-Filho et al., 2017; Yang et al., 2017).

After divisome assembly, cells constrict inward via envelope remodeling. In Gram-negative bacteria, the cell envelope is composed of an inner membrane, cell wall, and outer membrane external to the cell wall. The cell wall is a macromolecular meshwork made of peptidoglycan (PG) that protects against turgor pressure and other stresses, and dictates bacterial morphology (Daitch and Goley, 2020). New PG is synthesized by glycosyltransferases that polymerize lipid II, a lipid-linked disaccharide made of N-acetylmuramic acid and N-acetylglucosamine with a pentapeptide side chain. The peptide stems are crosslinked by transpeptidases, including D,D-transpeptidases of the penicillin-binding protein (PBP) family (Daitch and Goley, 2020). During division, the PG synthases FtsW and FtsI (PBP3) are the primary glycosyltransferase and transpeptidase, respectively (Daitch and Goley, 2020; Mahone and Goley, 2020). These two enzymes work together as a cognate pair (FtsWI) to synthesize cytokinetic PG that provides the constrictive force (Coltharp and Xiao, 2017).

Single molecule tracking studies show that the PG synthases move dynamically about the division site. In bacteria where dynamics have been characterized (*Escherichia coli, Bacillus subtilis, Staphylococcus aureus, Streptococcus pneumoniae*), the dynamics of FtsZ and the PG synthases are associated, but range from FtsWI requiring FtsZ treadmilling for movement to requiring FtsZ for placement at midcell, but moving independently (Yang et al., 2017, 2021; Bisson-Filho et al., 2017; Perez et al., 2019; Yang and Liu, 2022). In *E. coli,* the only Gram-negative organism in which dynamics have been studied, the moving PG synthases can be differentiated into fast- and slow-moving populations. Fast-moving PG synthases depend on FtsZ for movement and are inactive for PG synthesis. Slow-moving molecules depend on PG synthesis for locomotion and are thought to be actively synthesizing new PG (Yang et al., 2017, 2021; McCausland et al., 2021).

In most model bacteria, divisome components upstream of FtsWI activation have been identified, but their precise functions and mechanisms of signaling remain unclear. In *E. coli*, FtsN has been proposed as a trigger for constriction initiation and is last to localize to the division plane (Weiss, 2015; Mahone and Goley, 2020; Lyu et al., 2022). Hyperactivating mutations in *ftsB* and *ftsL*, which encode subunits of the FtsQLB complex, result in shorter cells (Tsang and Bernhardt, 2015; Liu et al., 2015). These mutants, when combined with other genetic perturbations, can bypass requirements of otherwise essential divisome components, suggesting they are critical mediators of activation (Park et al., 2020; Li et al., 2021; Du et al., 2016). Consistent with this idea, purified FtsQLB from *Pseudomonas aeruginosa* can activate FtsWI *in vitro* (Marmont and Bernhardt, 2020). While the final stages of activation are becoming clear, the FtsZ-proximal partners and how they ultimately signal for FtsWI activation remain elusive.

In *Caulobacter crescentus,* an α-proteobacterium in which divisome assembly has been well-defined, FtsW is the last divisome component to arrive at the division plane prior to constriction initiation (Goley, Yeh et al., 2011). However, localization of FtsW is not sufficient for cells to constrict. Recently, we implicated FzlA, an essential division protein conserved across α-proteobacteria, as a regulator of constriction (Lariviere et al., 2019, 2018; Goley, 2010). FzlA binds to FtsZ and co-localizes with FtsZ in stalked and pre-divisional cells (Goley, 2010). Previous work identified residues on FzlA required for two essential activities: binding to FtsZ and an unknown activity in the C-terminal tail (Lariviere et al., 2018). When FzlA is depleted, or either essential activity is disrupted, cells do not activate constriction, despite the remainder of the divisome localizing to midcell rings (Lariviere et al., 2018; Goley, 2010). A hyperactive mutant of FtsW (FtsW^A246T^) renders *fzlA* non-essential, albeit with a slower constriction rate than in the presence of FzlA (Lariviere et al., 2019). An *ftsWI* triple mutant (*ftsW^F145L/A246T^ftsI^I45V^,* termed *ftsW**I\**) constricts faster than the *ftsW^A246T^* mutant and also renders *fzlA* non-essential, with minimal effects on cell length (Lariviere et al., 2019; Lambert et al., 2018). We concluded that FzlA acts upstream of FtsWI activation, as an FtsZ-proximal step in the FtsZ-FtsWI activation pathway.

While we have established that FzlA is required for FtsWI activation in *Caulobacter*, the mechanism by which FzlA signals for FtsWI activation and the functional importance of regulation by FzlA is unknown. Here, we employed time-lapse imaging, genetics, biochemistry, and single-molecule tracking (SMT) to dissect the relationship between FzlA and FtsWI activity. We discovered that FzlA activates constriction by converting inactive, fast-moving molecules of FtsW to an active, slower-moving state. Overproducing FzlA leads to hyperconstriction by converting a greater proportion of FtsW into an active state, and this hyperactivation is toxic, particularly in the *ftsW**I\** background. The toxicity of FzlA overproduction is in part due to enhanced DNA damage via misregulation of a novel interaction between FzlA and FtsK, a division protein that segregates the chromosomal termini during division (Wang and Lutkenhaus, 1998; Wang et al., 2006). These results unify divisome assembly, chromosome segregation, and activation of cytokinetic PG synthases into a single pathway, which requires proper regulation to ensure envelope and chromosomal integrity.

## Results

### Overproduction of FzlA hyperactivates constriction

To better understand how FzlA regulates FtsWI, we sought to characterize the effects of FzlA overproduction on cell division and viability. We hypothesized that FzlA may be a limiting activator of FtsWI. If that is the case, cells with an overabundance of FzlA should be hyperactive for constriction and therefore constrict faster than control cells (Yang et al., 2021; Lambert et al., 2018; Lariviere et al., 2019). To test this hypothesis, we performed timelapse phase contrast microscopy on synchronized cells overexpressing *fzlA* (EG3637) or bearing an empty vector (EV) control (EG1644) (Fig. 1 A). We induced *fzlA* overexpression for one hour prior to and throughout the timelapse experiments, and isolated newborn swarmer (G1) cells by density centrifugation prior to imaging. We determined that *fzlA-*overexpressing cells increase FzlA levels to roughly 20-fold higher than WT, as measured by western blotting with an anti-FzlA antibody (Fig. S1, A-B). For each cell, we calculated the rates of constriction and elongation as well as the time from birth to initiation of constriction (pre-constriction time).

**Figure 1.**
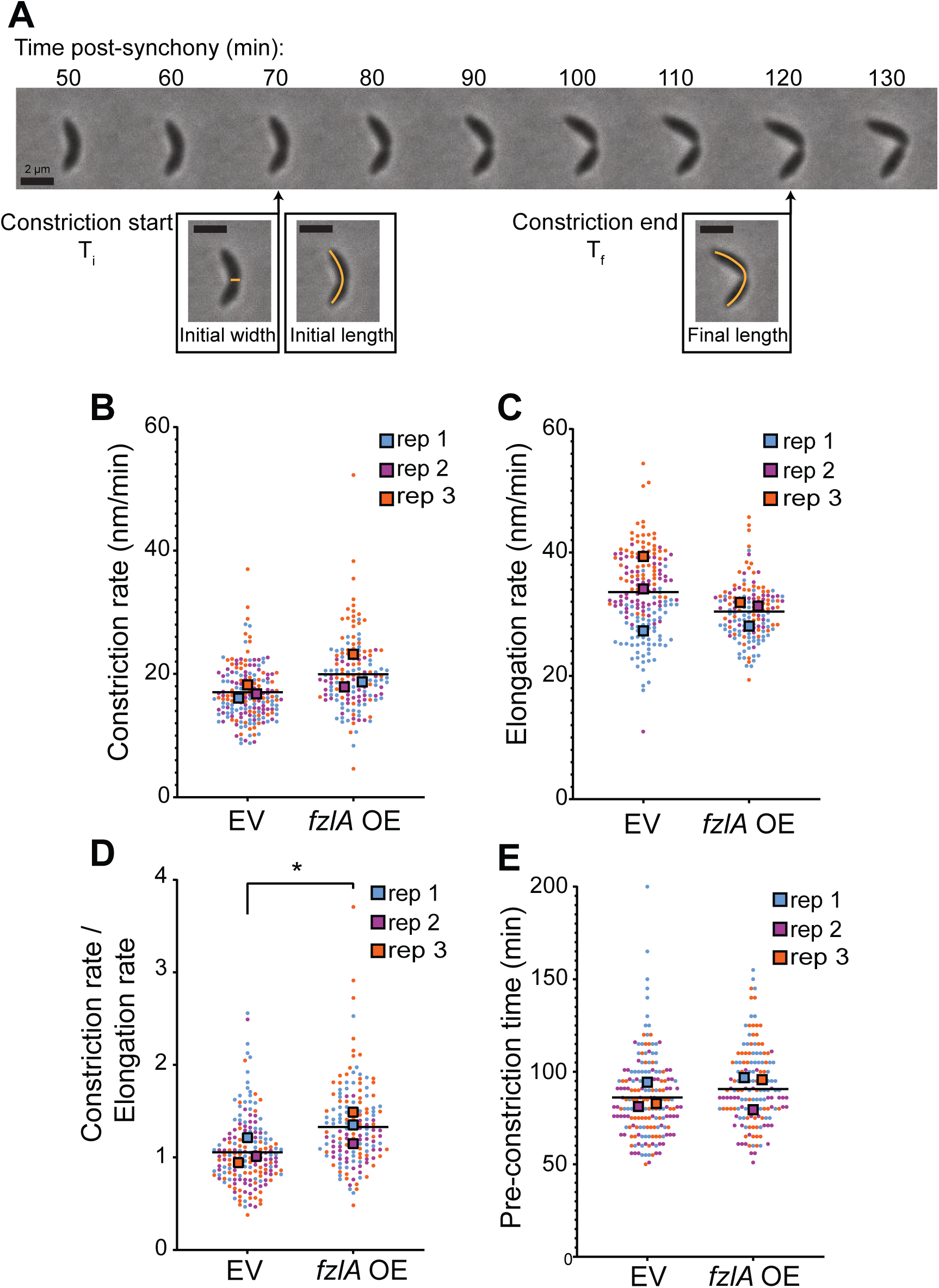
Overproduction of FzlA hyperactivates constriction. **A.** Representative phase contrast timelapse microscopy of a wild-type constricting *C. crescentus* cell. The timepoints of constriction initiation and completion are denoted. Scale bars are all 2 µm. The orange lines are drawn for ease of visualizing the approximate measures of length or width for each cell. T_i_: Time of initiation. T_f_: Final timepoint of constriction. **B-E.** Superplots comparing the **B.** constriction rate **C.** elongation rate **D.** ratio of constriction to elongation rate or **E.** time to constriction initiation for strains harboring either an empty vector (EV; EG1644) control or *fzlA* overexpression (OE; EG3637) construct. The strains were induced with 0.3% xylose for 1 h prior to and throughout the timelapse experiment. Each circle is the value of a single cell, while the large squares are the average of a biological replicate. The bar is the average of the three means and statistics were performed using the means of the three replicates. EG1644 – replicate 1, 67 cells; replicate 2, 63 cells; replicate 3, 54 cells. EG3637 – replicate 1, 60 cells; replicate 2, 40 cells; replicate 3, 55 cells. * represents a p-value < 0.05.

When FzlA was overproduced, cells constricted significantly faster on average (20.0 ± 0.5 nm/min, N = 154) than the EV control (16.8 ± 0.3 nm/min, N = 182) using conventional statistical analysis (Fig. 1B, Video 1, p < 0.0001). To ensure that the measured differences were not due to sample size effects, we performed a more strenuous analysis based on Superplots, a method of statistical analysis that treats the averages of independent biological replicate experiments (rather than individual cell values) as data points for each condition (Lord et al., 2020). With Superplots analysis, the constriction rate was still significantly faster when *fzlA* was over-expressed (19.8 ± 1.6 nm/min) compared to an EV (16.9 ± 0.6 nm/min), using an α-cutoff of 0.1, but not when using an α-cutoff of 0.05 (Fig. 1B, p = 0.1). In *Caulobacter*, the elongation rate is generally inversely proportional to the constriction rate. That is, as cells constrict faster, they elongate slower and vice versa (Lambert et al., 2018; Lariviere et al., 2019). Superplots analysis of the elongation rates showed no significant difference between *fzlA* over-expressing (15.2 ± 0.6 nm/min) and EV control cells (16.8 ± 1.7 nm/min) (Fig. 1C). In contrast, the ratio of constriction rate to elongation rate by Superplots analysis was significantly shifted in favor of constriction in strains overproducing FzlA (1.33 ± 0.10) when compared to EV control cells (1.06 ± 0.08), signifying a shift from elongation to constriction, similar to observations made for hyperactive strains (Fig. 1D, p = 0.049) (Lariviere et al., 2019). These results demonstrate that *fzlA* overexpression enhances constriction, consistent with the hypothesis that FzlA is a limiting factor in FtsWI activation. We used the amount of time elapsed post-synchrony to measure the time from birth to constriction initiation. There was no significant difference in time to constriction initiation between *fzlA* overexpressing (92 ± 2 min, N = 154) and EV control cells (86 ± 2 min, N = 182), even by conventional statistics (Fig. 1E, p = 0.071), indicating that excess FzlA does not pre-maturely initiate constriction, but rather causes FtsWI to build PG inwards at an increased rate.

### Active Halo-FtsW molecules move more slowly on average

We hypothesized that *fzlA* overexpressing cells constrict faster because FtsWI are synthesizing PG more frequently or at an increased rate. To test these hypotheses, we performed single molecule tracking (SMT) of FtsW. In *E. coli,* the phylogenetically closest relative in which PG synthase dynamics have been measured, inactive PG synthases move with similar speeds (32 nm/s) to the FtsZ-treadmilling speed (28 nm/s) but move more slowly (8 nm/s) when activated (Yang et al., 2021). We therefore sought to characterize the relationship between FtsWI activation state and dynamics in *C. crescentus* by performing SMT.

We generated strains expressing either *halo-ftsW or -ftsW\*\** as the sole copy at the native locus, in a WT (EG3052) or *ftsW**I\** (EG3053) background, respectively. We labeled and measured single molecules of FtsW or FtsW** by titrating the levels of Janelia Fluor 646 Halo ligand (Grimm et al., 2017). While tracking FtsW, we observed single molecules that had processive movement, with some molecules changing direction and speed, implying that FtsW can dynamically switch between movement modes, similar to what was observed in *E. coli* (Yang et al., 2021). To focus on FtsW molecules during division, we synchronized cells prior to analysis and imaged after approximately 50 minutes of growth to ensure we were imaging pre-divisional and constricting cells. We measured speeds of mobile molecules of FtsW at the division site by dividing the distances covered by the time each molecule took to cover that distance (Fig. 2A, Fig. S2, A-B, Video 2-4). In some cases, a single molecule exhibited changes in the speed of movement and/or transitioned from moving to stationary or vice versa during measurement (Fig. S2, A-B, Video 3-4). Additionally, a proportion of molecules existed at the division plane without movement, which we define as the stationary population. We measured the number of molecules in a population that were stationary, as well as how long they remained stationary.

**Figure 2.**
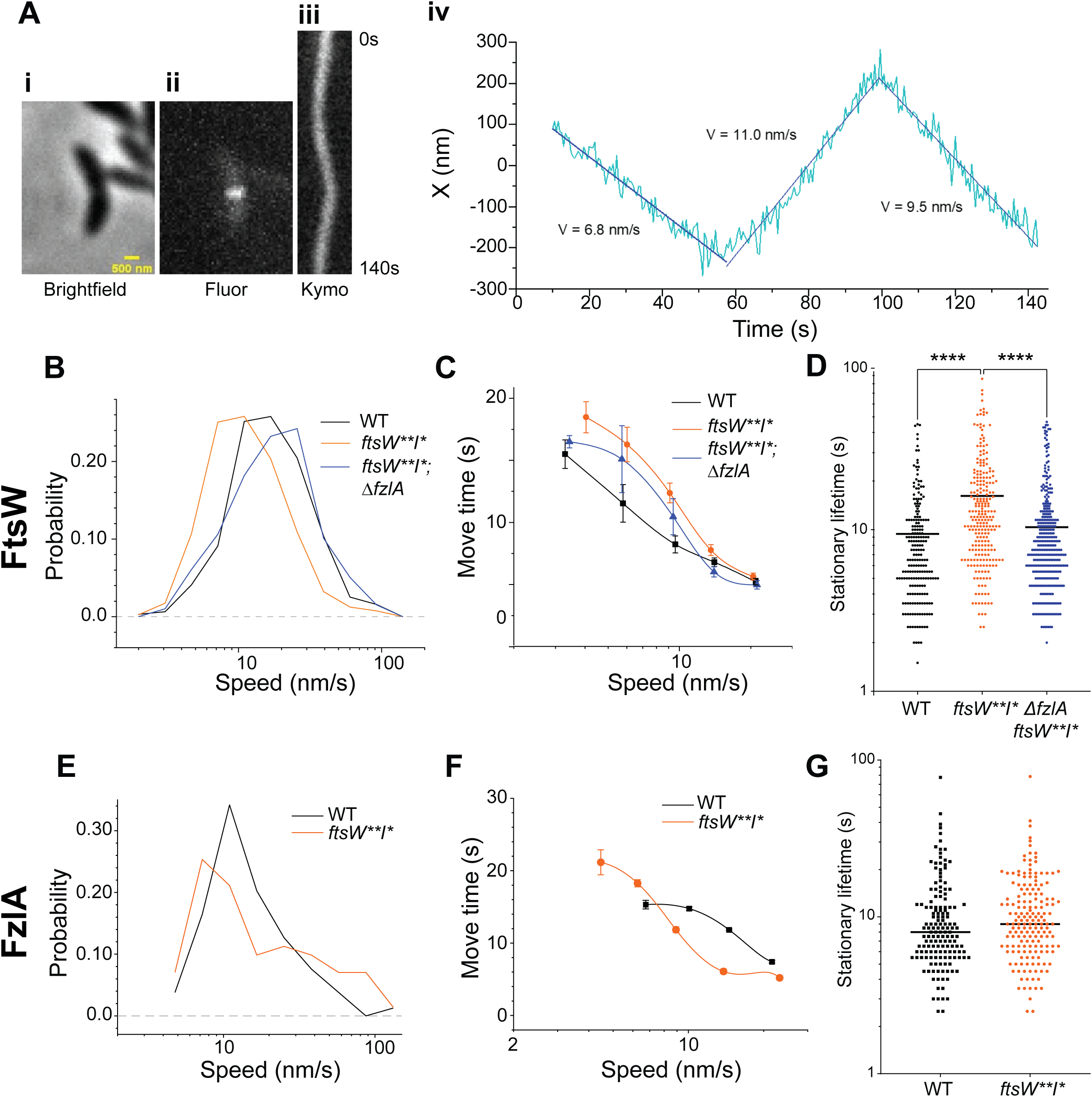
FtsW and FzlA influence each other’s movement. **A. i.** Brightfield image of a *Halo-ftsW**I\** cell. **ii.** Representative maximum fluorescence intensity projection (Fluor) for Halo-FtsW**. **iii.** Kymograph (Kymo) of the fluorescence signal of a line scan across the division plane for a Janelia Fluor 646-labeled single Halo-FtsW**. **iv.** Plot of the molecule position within the line scan over the course of timelapse imaging and speeds of movement for each segment. X indicates the short axis of the cell. **B.** Histogram comparing single molecule speeds of FtsW or FtsW** in *halo-ftsW* (WT; EG3052, N = 318), *halo-ftsW\*\**; *ftsI\** (*ftsW**I\**; EG3053, N = 403), or *ΔfzlA*; *halo-ftsW\*\**; *ftsI\** (*ΔfzlA, ftsW**I\**; EG3207, N = 198) backgrounds. **C.** Plot comparing the lifetime of single FtsW or FtsW** molecules move at slow speeds (< 20 nm/s) in *halo-ftsW* (WT; EG3052, N = 318), *halo-ftsW\*\**; *ftsI\** (*ftsW**I\**; EG3053, N = 403), or *ΔfzlA*; *halo-ftsW\*\**, *ftsI\** (*ΔfzlA, ftsW**I\**; EG3207, N = 198) backgrounds. Error bars are the standard error of the mean for the speeds in the move-times of those molecules. **D.** Plot comparing the lifetimes of stationary single FtsW or FtsW** molecules in a *halo-ftsW* (WT; EG3052, N = 210), *halo-ftsW\*\**; *ftsI\** (*ftsW**I\**; EG3053, N = 249), or *ΔfzlA*; *halo-ftsW\*\**; *ftsI\** (*ΔfzlA, ftsW**I\**; EG3207, N = 366) backgrounds. **E.** Histogram comparing speeds of FzlA molecules in *halo-fzlA*; WT (EG3619, N = 104) or *halo-fzlA*; *ftsW**I\** (EG3617, N = 71) backgrounds. **F.** Plot comparing the length of time single FzlA molecules move at slow speeds (< 20 nm/s) in *halo-fzlA;* WT (EG3619, N = 318) or *halo-fzlA; ftsW**I\** (EG3617, N = 403) backgrounds. Error bars are the standard error of the mean for the speeds in the move-times of those molecules. **G.** Plot comparing the lifetimes of stationary FzlA molecules in a *halo-fzlA;* WT (EG3619, 153 lifetimes) or *halo-fzlA*; *ftsW**I\** (EG3617, 174 lifetimes) background. **** represents a p-value < 0.0001.

We observed that hyperactivated FtsW** molecules move more slowly on average (16.6 ± 0.7 nm/s, N = 403) than WT FtsW (24.9 ± 1.1 nm/s, N = 318) (Fig. 2B, Table 1), consistent with observations in *E. coli* that active FtsWI molecules move at a slower rate than inactive, but mobile, molecules in *E. coli* (Yang et al., 2021). To compare the processivity of mobile FtsW molecules between the *ftsW**I\** and WT backgrounds, we plotted the speeds of moving molecules against the time those molecules spent on moving. For this analysis, we excluded molecules moving faster than 20 nm/s as they are unlikely to be actively synthesizing PG. We found that slow-moving FtsW** molecules travel for longer periods of time than slow-moving WT FtsW on average (Fig. 2C). Since slow-moving PG synthases are more likely to be active, these data suggest active FtsW** molecules are more processive than active WT FtsW molecules. Interestingly, we did not observe a change in the percentage of stationary FtsW molecules when FtsW was hyperactivated (roughly 50% in both WT and *ftsW**I\**), but instead observed longer stationary lifetimes for FtsW** molecules (16.2 ± 0.8 s, N = 249) when compared to WT (9.4 ± 0.5 s, N = 210) (Fig. 2D, Table 1). Based on observations in *E. coli*, we hypothesize that stationary molecules are not active for cell wall synthesis but may be poised for activation (Yang et al., 2021). Our data indicate that *ftsW**I\** cells constrict faster due to both a greater proportion of active (slow moving) PG synthase complexes and because these complexes polymerize longer stretches of cytokinetic PG than their WT counterparts.

### Absence of FzlA decreases the proportion of active Halo-FtsW** in a *ftsW**I\** background

To understand the role FzlA plays in FtsWI activation, we deleted *fzlA* in the *ftsW**I\** background producing Halo-FtsW** and replaced it with a gentamycin resistance cassette, confirmed by western blotting (Fig. S1 C). Notably, in the absence of FzlA (EG3207), moving FtsW** molecules (26.1 ± 1.6 nm/s, N = 198) had average speeds similar to FtsW in a WT background (24.9 ± 1.1 nm/s, N = 318) and significantly faster than FtsW** in the presence of FzlA (16.6 ± 0.7, N = 403) (Table 1, Fig. 2B). However, the proportion of stationary PG synthases significantly increased when compared to either FtsW** or FtsW in the presence of FzlA (Table S1, 75% in *ΔfzlA* vs ∼50% in WT or *ftsW**I\**). These data suggest that there are significantly fewer active, slow-moving synthases in a *ΔfzlA* background than WT or *ftsW**I\** with FzlA and are consistent with the cell length and constriction rate defects associated with *fzlA* deletion we previously reported (Lariviere et al., 2019). These data further indicate that FtsW**I* can still receive input from FzlA even though *fzlA* is no longer essential.

Interestingly, FtsW** in the *ΔfzlA* background appeared to have processive movement similar to FtsW** in the presence of FzlA, suggesting that FzlA does not affect the processivity of PG synthases, but rather the fraction of active PG synthases (Fig. 2C). Finally, absence of FzlA reduced the amount of time FtsW** molecules remained stationary when compared to stationary time in the presence of FzlA (Table S1, Fig. 2D, 10.4 ± 0.4 s, N = 366 for *ΔfzlA; ftsW**I\** vs 16.2 ± 0.8 s, N = 249 for *ftsW**I\**), to an average stationary time similar to WT FtsW (Table S1, Fig. 2D, 9.4 ± 0.5 s, N = 210). Since the absence of FzlA increases the stationary proportion of FtsW** molecules and these molecules are shorter-lived than when FzlA is present, we propose that short-lived stationary FtsW are poised, awaiting activation by a FzlA-dependent pathway.

### FzlA moves with active FtsWI

Considering that FzlA is an FtsWI activator, we sought to test if FzlA moves with activated FtsWI or functions through a “kiss-and-run” mechanism, dissociating shortly after activation. To do this, we assessed FzlA dynamics using SMT. We constructed strains expressing *halo-fzlA* at the native *fzlA* locus in WT (EG3619) and *ftsW**I\** (EG3617) backgrounds. We observed similar average FzlA speeds in the WT (Table S1, 17.3 ± 1.1 nm/s, N = 104) and *ftsW**I\** backgrounds (Table S1, 20.3 ± 2.5 nm/s, N = 71). However, the distribution of Halo-FzlA speeds was broader in a *ftsW**I\** background than in WT, with more very slow- and very fast-moving molecules than in WT (Fig. 2E). The presence of slow-moving FzlA molecules suggests that it can remain associated with activated FtsWI as it synthesizes PG. In the *ftsW**I\** background, FzlA was stationary more often than in WT cells (Table S1, 72% stationary in *ftsW**I\** vs 59% stationary in WT). The activation state of the PG synthases impacted the slow-moving population of FzlA (Fig. 2F), but not the stationary life-time of FzlA (Fig. 2G), leading us to conclude that FzlA likely remains associated with active, slow-moving FtsWI complexes after activation.

### Absence of FzlA does not impact the proportion of stationary Halo-FtsW in a WT background

To examine the effects of loss of FzlA on FtsW dynamics in a WT background, we generated a strain dependent on xylose for *fzlA* expression with natively expressed WT *ftsWI* (EG3523). When the xylose inducer was removed, FzlA dropped to undetectable levels after seven hours and cells began to filament (Fig. S3, Fig. S1 D). We optimized xylose concentration to produce FzlA at WT levels and avoid FtsWI hyperactivation by overexpression of *fzlA* before depletion (Fig. S1 E). To ensure that we measured FtsW molecules that are associated with a Z-ring in these filamentous cells, we replaced *zapA* at the native locus with *zapA-mNeonGreen* (*zapA-mNG*) (Fig. S3). ZapA is an FtsZ-binding protein that reliably reports on Z-ring positioning (Woldemeskel et al., 2017).

When comparing FtsW dynamics between induced and depleted conditions, we observed that FtsW moves faster, on average, when FzlA is absent (Table S1, Fig. 3A, 23.0 ± 1.9 nm/s during FzlA depletion, N = 117 vs 19.6 ± 1.1 nm/s with FzlA present, N = 230), consistent with our *ΔfzlA* results and suggesting fewer active PG synthase complexes are present without FzlA. FtsW processivity was not affected by FzlA in this background (Fig. 3B), consistent with our observations using the *ftsW**I\** strain with and without FzlA.

**Figure 3.**
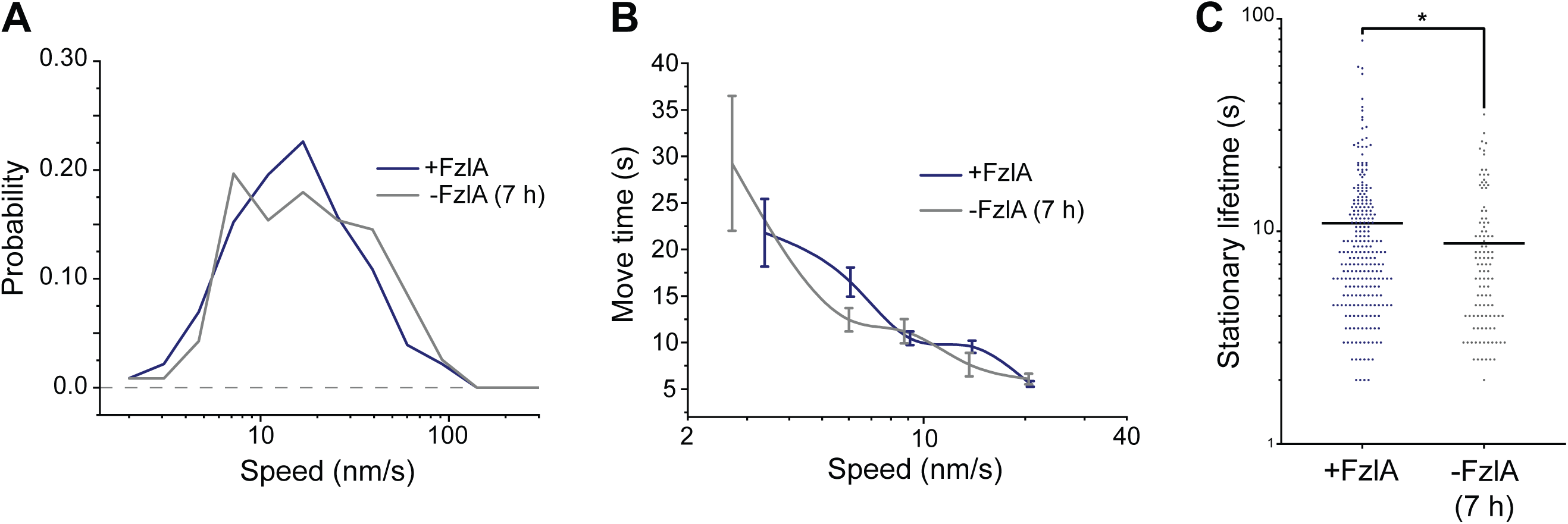
Depletion of FzlA increases average FtsW speeds and decreases stationary FtsW lifetimes in a WT background. **A.** Histogram comparing single molecule speeds of FtsW in cells (EG3523) producing FzlA (+FzlA, 0.001% xylose, N = 230) or depleted of FzlA for seven hours (-FzlA (7 h), 0.001% glucose, N = 117). **B.** Plot comparing the lifetime of single FtsW molecules moving at slow speeds (< 20 nm/s) in cells producing FzlA (+FzlA, 0.001% xylose, N = 230) or depleted of FzlA for seven hours (-FzlA (7 h), 0.001% glucose, N = 117). **C.** Plot comparing the lifetimes of stationary single FtsW molecules in cells producing FzlA (+FzlA, 0.001% xylose, N = 244) or depleted of FzlA for seven hours (-FzlA (7 h), 0.001% glucose, N = 107). * represents a p-value < 0.05.

We next assessed the stationary FtsW population in our depletion strain with and without FzlA. Surprisingly, depletion of FzlA in a WT background did not change the proportion of moving FtsW molecules (Table S1, ∼50% in both deplete and induced), unlike in the *ΔfzlA* background (Table S1, 25%). These results indicate that the increase in stationary FtsW** molecules observed in the *ΔfzlA* background is likely due to binding of the PG synthases to a stationary target rather than an inability to move dynamically about the Z-ring in the absence of FzlA. The amount of time FtsW molecules remain stationary decreased when FzlA was depleted in a WT background (Table S1, Fig. 3C, 8.8 ± 0.7 s during FzlA depletion, N = 107 vs 10.9 ± 0.6 s at WT FzlA levels, N = 244). These tracking data are consistent with FzlA-mediated signaling acting to convert fast-moving, inactive molecules of FtsW into an activated, slow-moving state. Moreover, our data indicate that FtsW molecules may become stationary when they are waiting to receive activating signals (i.e., via FzlA) or when they are poised to synthesize PG but have not yet begun to do so (i.e., for FtsW**I*).

### Overabundant FzlA converts Halo-FtsW from a fast-moving and inactive state to a slow and active PG-synthesis state

Having assessed the effects of loss of FzlA on FtsWI dynamics and activity, we next turned to cells overproducing FzlA. To identify changes to PG synthase activation during hyperconstriction caused by FzlA overproduction, we compared Halo-FtsW speeds from strains with the *fzlA* overexpression construct (EG3519) or EV (EG3537) in the presence of 0.3% xylose to induce *fzlA* overexpression for one hour prior to synchrony and during the experiment (Fig. S1 F). With FzlA overproduction, FtsW speeds were reduced compared to EV (Table S1, Fig. 4A, 23.2 ± 0.9 nm/s with FzlA overproduction, N = 300 vs 32.3 ± 3.6 nm/s for the EV control, N = 472). As expected from earlier results, additional FzlA altered neither the processivity nor the proportion of moving molecules (Table S1, Fig. 4B). These results reinforce our conclusion that FzlA-mediated signaling converts inactive PG synthases into an active state.

**Figure 4.**
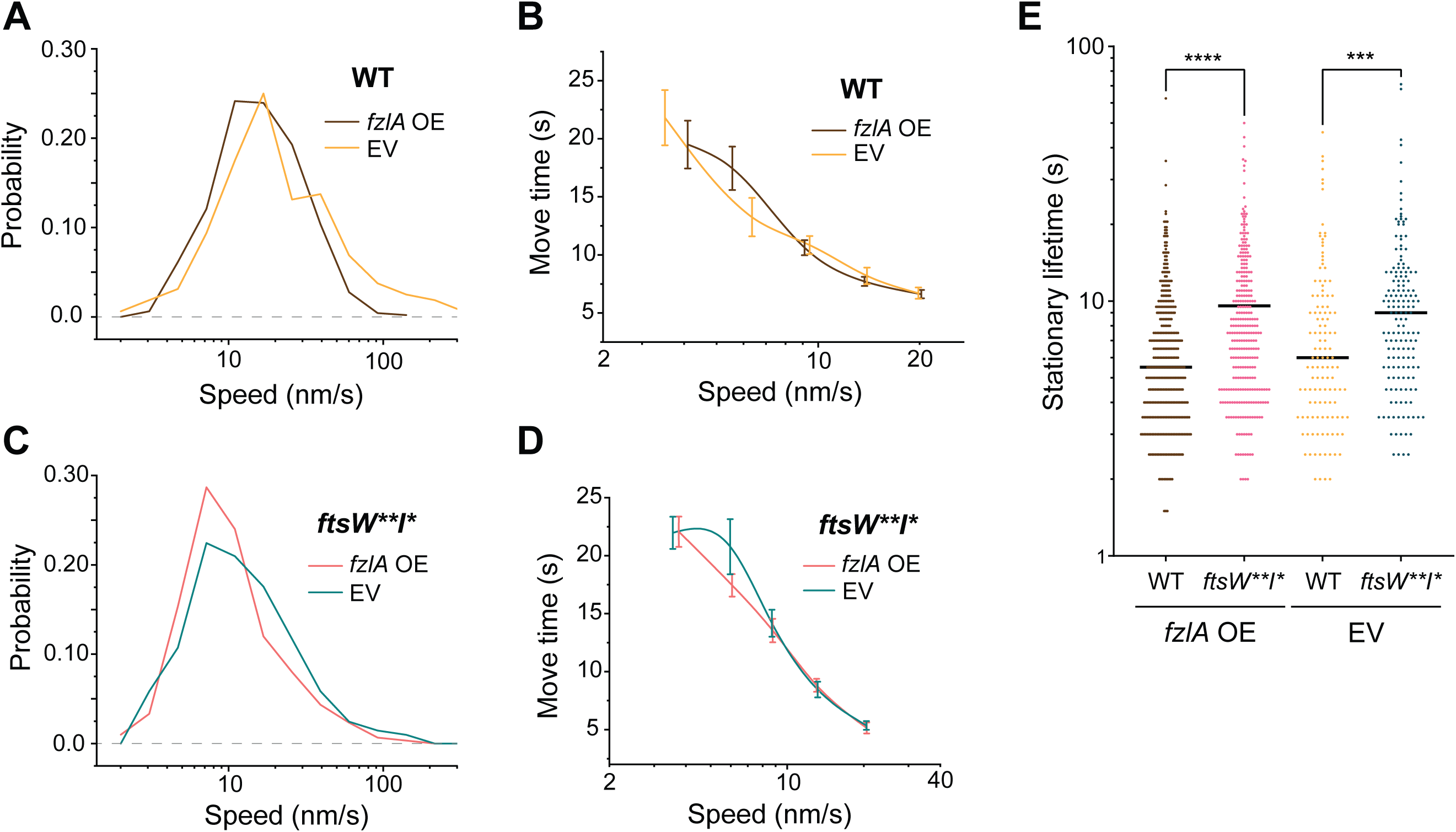
FtsW and FtsW** molecules move more slowly when FzlA is overabundant. **A and C.** Histogram comparing speeds of FtsW or FtsW** in cells with an empty vector (EV) control or with a *fzlA* overexpression (OE) construct in either **A.** a *halo-ftsW* (EV; EG3537, N = 472. *fzlA* OE; EG3519, N = 300) or **C.** *halo-ftsW\*\**; *ftsI\** (EV; EG3538, N = 160. *fzlA* OE; EG3525, N = 205) background. **B and D.** Plot comparing the lifetime of single FtsW and FtsW** molecules moving at slow speeds (< 20 nm/s) with and without FzlA overproduction in a **B.** *halo-ftsW* or **D.** *halo-ftsW\*\**; *ftsI\** background. **E.** Plot of the lifetimes of stationary single FtsW or FtsW** molecules during *fzlA* overexpression (OE) compared to an empty vector (EV) control in a *halo-ftsW* or *halo-ftsW\*\**; *ftsI\** background (WT; *fzlA* OE, EG3519, N = 291. *ftsW**I\**; *fzlA* OE, EG3525, N = 159. WT; EV, EG3537, N = 381. EG3538, N = 123). 0.3% xylose was used to overexpress *fzlA*. WT: wild-type apart from *halo-ftsW*. **** represents a p-value < 0.0001. *** represents a p-value < 0.001.

Our prior results demonstrated that FzlA-mediated signaling still impacts FtsW**, which suggests that FzlA-mediated hyperactivation can occur in a *ftsW**I\** background as well. In line with this hypothesis, we observed that FzlA overproduction in this background (EG3525) further stimulates FtsW**, as speeds decreased to the slowest observed average speed when FzlA was overproduced compared to EV (EG3538) (Table S1, Fig. 4C, 13.5 ± 0.8 nm/s for overproduced FzlA, N = 205 vs 16.3 ± 1.2 nm/s for EV, N = 160). Again, there was no change to processivity or the proportion of moving PG synthases when FzlA was overproduced (Table S1, Fig. 4D). These results further demonstrate that in the *ftsW**I\** background, the PG synthases are primed for activation, but FtsW**I* can still receive activation signals from FzlA. However, excess FzlA again had no effect on the proportion of stationary FtsW or FtsW** or their lifetimes (Table S1, Fig. 4E).

### FtsW and FzlA molecules exhibit two moving populations

In *E. coli*, the PG synthases move with two dynamic modes: 1) fast, inactive PG synthases driven by FtsZ cluster movement and 2) slow, actively synthesizing PG synthases driven by synthesis (Yang et al., 2021). Due to the small size of *C. crescentus*, we were unable to resolve FtsZ clusters sufficiently to determine their treadmilling speeds. To better understand PG synthase dynamics in *C. crescentus*, we employed a computational approach to simulate the contributions of two populations of moving molecules to the distributions of FtsW and FzlA speeds we observed. In a WT background, FtsW speeds appear to move as a major population with an average speed of 22.2 nm/s and a small portion moving much faster at an average speed of 54.6 nm/s (Fig. 5Ai, Table S1), although the two-populations were not well-distinguished. However, when observing FtsW** in the hyperactive background, slow- (Fig. 5Aii, Table S1, 10.8 ± 1.1 nm/s) and fast-moving (Fig. 5Aii, Table S1, 51.4 ± 11.4 nm/s) populations are better resolved, suggesting that there are two-populations of FtsW and that the actively synthesizing population moves slower on average. To understand if FzlA modulates FtsW dynamics by shifting FtsW from one population to the other, we compared the proportions of each FtsW population in *ftsW**I\** backgrounds with and without FzlA. In a *ΔfzlA* background, a greater proportion of FtsW moved at faster speeds (Fig. 5Aiii, Table S1, 84.2% fast-moving) than when FzlA was present (Fig. 5Aiii, Table S1, 17.1% fast-moving), suggesting less active FtsW when FzlA was not present. This trend was observed when FzlA was not present in a WT background as well, and the inverse was observed when FzlA was overproduced, solidifying the ability of FzlA to convert fast-moving FtsW to a slow-moving and active state (Fig. S4, Table S1).

**Figure 5.**
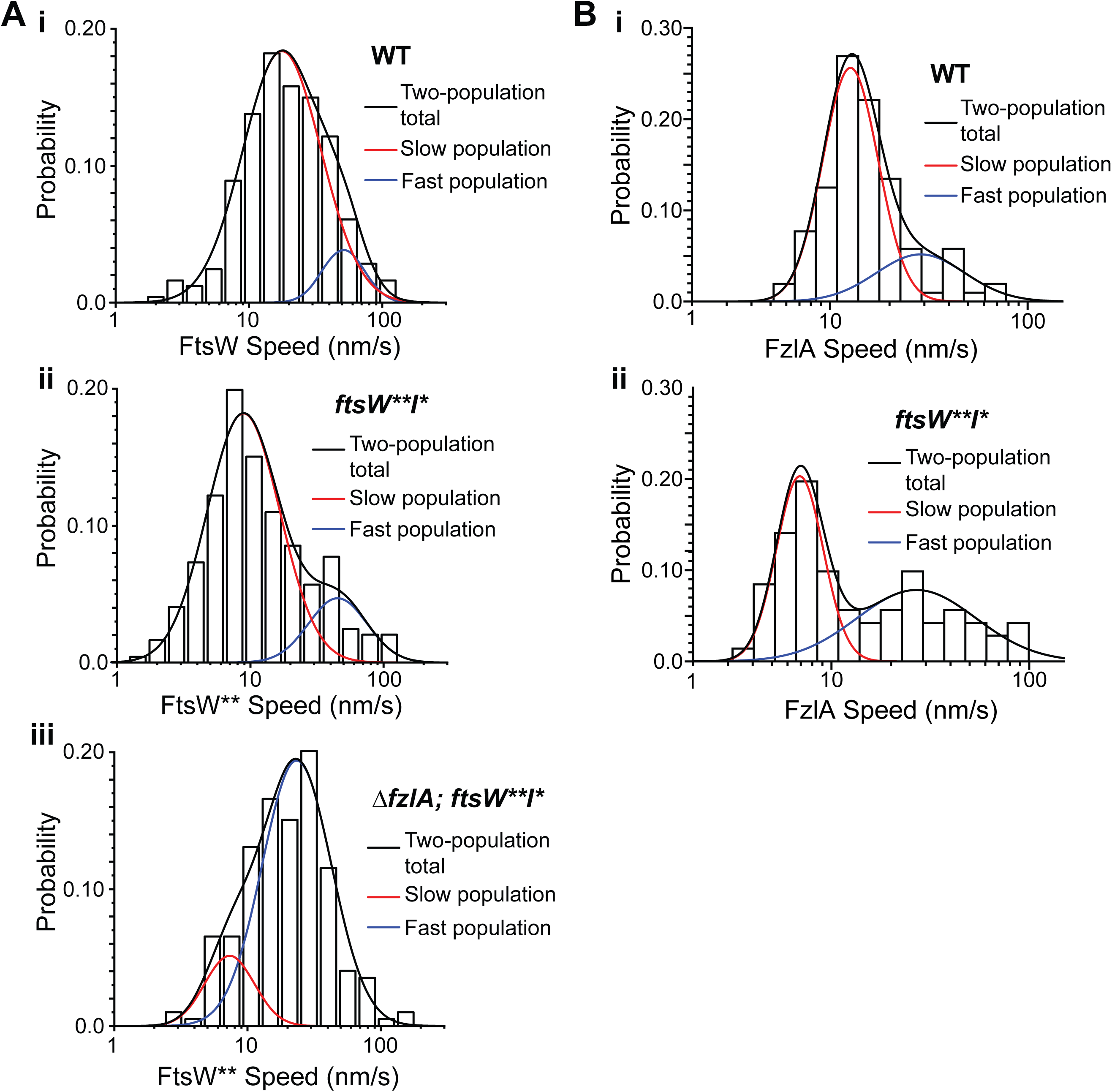
Active FtsW moves at a slower rate than inactive FtsW, while FzlA moves with active FtsW. **A-D.** Histograms of SMT data for Halo fusion proteins with calculated two-population curves that best fit the data for **A.** FtsW or FtsW** in a **i.** *halo-ftsW* (WT; EG3052), **i.** *halo-ftsW\*\**; *ftsI\** (*ftsW**I\**; EG3053), or **iii.** *ΔfzlA*; *halo-ftsW\*\**; *ftsI\** (*ΔfzlA, ftsW**I\**; EG3207) background (a re-analysis of the data presented in Fig. 2B). **B.** FzlA in **i.** a WT (EG3619) and **ii.** *ftsW**I\** (EG3617) background (a re-analysis of the data presented in Fig. 2E). Red = slow population. Blue = fast population. Black: total based on two-population.

After confirming FzlA can shift FtsW from an inactive, fast-moving state to a slow-moving active state, we sought to understand the effects of FtsW activation state on FzlA dynamics. In the WT background, FzlA appears to move primarily in a slow population (Fig. 5Bi, Table S1, 13.4 ± 1.0 nm/s, 76% slow-moving FzlA). When FtsW is hyperactivated, the slow-moving population was observed moving even slower on average (Fig. 5Bii, Table S1, 7.2 ± 0.5 nm/s). This is consistent with our hypothesis that moving FzlA travels with active FtsW (Fig. 5Ai and ii).

### Hyperactivated constriction via FzlA overproduction causes lethal division events

Previous work demonstrated that hyperactivating FtsWI has consequences on the cell wall and envelope integrity (Modell et al., 2014; Lariviere et al., 2019). These observations led us to ask if there are phenotypic consequences of FtsWI hyperactivation through an over-abundance of FzlA. While analyzing timelapse images of *fzlA* overexpressing cells, we noticed that cells died during or immediately after division at a significant rate, though a majority of cells were still viable. The lethal division events we observed were pleiotropic, with three major distinguishable categories; 1) both daughters halt growth post-division, 2) one daughter halts growth post-division, and 3) one daughter lyses post-division (Fig. 6, A-B, Video 5-7). These lethal division events were rare in the EV control strain, suggesting that the hyperconstriction caused by FzlA over-abundance is deleterious to cells.

**Figure 6.**
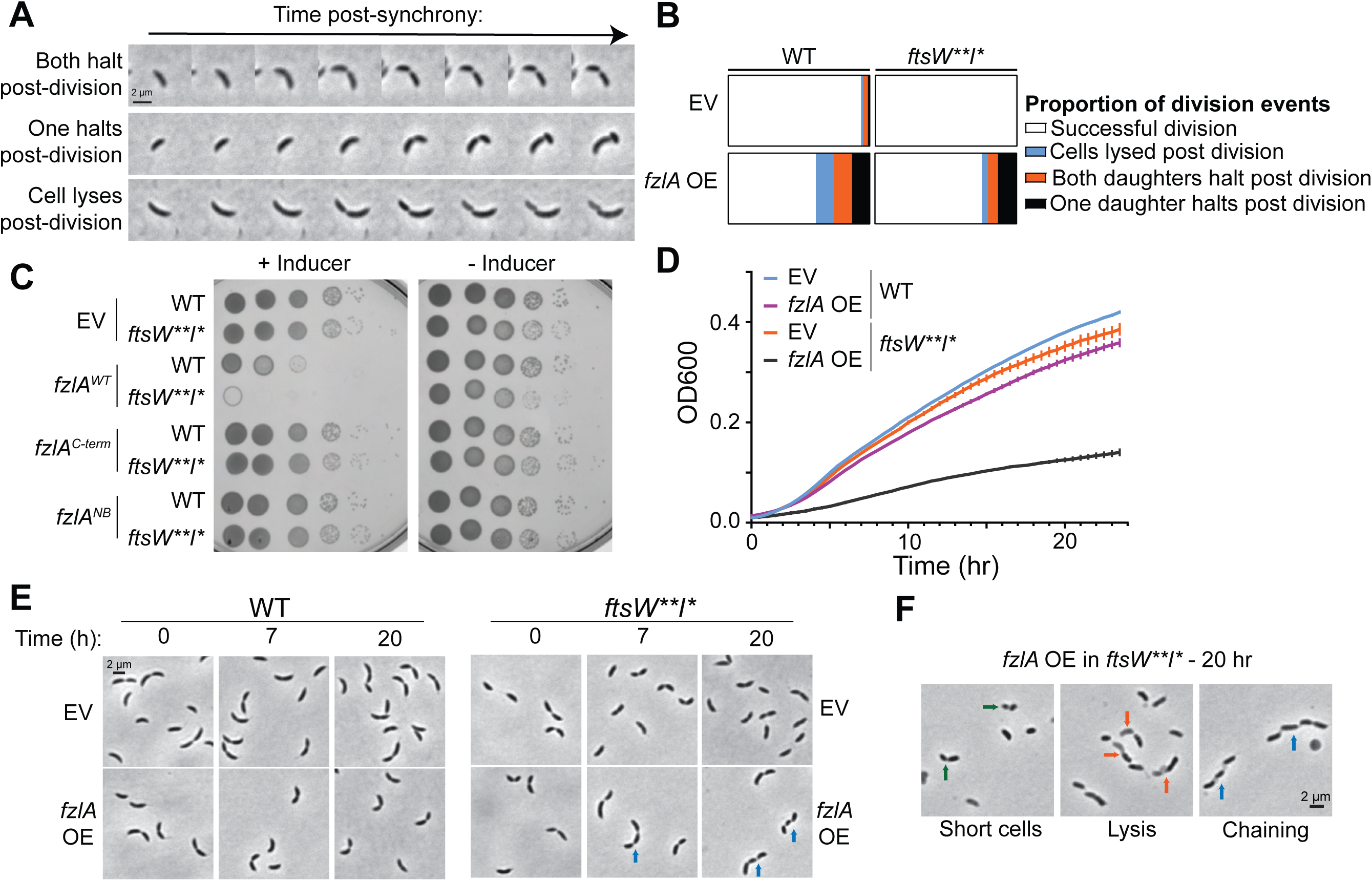
Overproduction of FzlA is toxic. **A.** Representative phase contrast timelapse images of constricting cells that result in lethal division events due to FzlA overproduction. **B.** Quantification of lethal divisions in the wild-type (WT) and *ftsW**I\** background comparing empty vector (EV) controls (WT: EG1644, n = 182. *ftsW**I\**: EG3466, n = 145) and to *fzlA* over-expression (WT: EG3637, n = 155. *ftsW**I\**: EG3467, n = 140). **C.** Spot dilutions of wild-type (WT) or *ftsW**I\** strains harboring an empty vector (EV) control or *fzlA* variant overexpression construct, plated with 0.3% xylose (+ Inducer) or 0.2% glucose (- Inducer). Cultures were diluted as indicated. **D.** Growth curves of wild-type (WT) or *ftsW**I\** strains harboring an empty vector (EV) control or *fzlA* overexpression construct. Strains were pre-induced for six hours before growth curves with 0.3% xylose and continued to be cultured with 0.3% xylose during the growth curve. Points are the mean of three technical replicates at that timepoint, error bars are the standard error of the mean. Shown is a representative replicate of a biological triplicate. **E.** Phase contrast time course (0, 7, 20 hr post-induction) of wild-type (WT) or *ftsW**I\** cells harboring an empty vector (EV) control or *fzlA* overexpression construct. **F.** Example phase contrast images of cells after 20 hours of FzlA overproduction in a *ftsW**I\** background. Arrow colors: Green = very short, Orange = one half of the constricted cell is lysed, Blue = cell chaining. *fzlA* OE, roughly 20-fold overproduced FzlA.

Interestingly, without FzlA overproduction in a *ftsW**I\** background (EG3466), we did not observe lethal divisions, suggesting hyperactive PG synthases are not sufficient to cause a significant increase in lethal division events on this timescale. FzlA overproduction in *ftsW**I\** (EG3467) increased lethal division events compared to the EV control, but not to a greater extent than in a WT background. To understand how these division failures affect long-term viability, we performed spot dilutions and growth curves in both WT and *ftsW**I\** backgrounds with and without FzlA overproduction. Overexpression of *fzlA* in a WT background caused a 2-log reduction in colony-forming units and a moderate reduction in doubling time in liquid media (Fig. 6, C-D). Strikingly, long-term overexpression of *fzlA* was lethal in a *ftsW**I\** background, both on solid and liquid medium (Fig. 6, C-D).

FzlA has two essential surfaces: an FtsZ-binding surface and the C-terminal tail (Lariviere et al., 2018). We next sought to test whether either function is required for the observed toxicity. To do this, we assessed the effects of overexpression of a mutant *fzlA* with a disrupted FtsZ-binding site (*fzlA^NB2^*) or a charge-reversal mutation in the penultimate residue of the C-terminal tail (*fzlA^D227K^)* in the WT or *ftsW**I\** backgrounds. Neither *fzlA^D227K^* nor *fzlA^NB2^* overexpression was lethal in WT or *ftsW**I\** backgrounds, demonstrating that FzlA must retain both essential activities for toxicity (Fig. 6C, Fig. S1, A-B).

To better understand how FzlA overproduction in a *ftsW**I\** background causes toxicity, we imaged cells after 20 hours of overproduction of FzlA. We observed cells that had lysed during division and remained attached (red arrows), similar to observations from the earlier time-lapse imaging, as well as cells that were significantly shorter (green arrows) than *ftsW**I\** cells with an EV (Fig. 6, E-F). Interestingly, we also observed cell chaining (blue arrows). The cells that chained frequently had multiple deep constrictions with cell bodies remaining connected, suggesting that division failed to complete at a late stage then initiated again at a second site.

### FtsK is a potential binding partner of FzlA

Cell chaining has been observed in cells with mutations to cell wall hydrolase-related proteins, outer-membrane components, and factors that regulate genome integrity and segregation (Meier et al., 2017; Bernhardt and de Boer, 2004; Peters et al., 2011; Heidrich et al., 2001; Wang et al., 2006; Uehara et al., 2010; lo Sciuto et al., 2014). To better understand both the mechanism for the cell division defects that result from FzlA overproduction and how FzlA signals to activate FtsWI, we sought to identify downstream interactors of FzlA. To identify FzlA binding partners, we performed co-immunoprecipitation (coIP) in a strain (EG2217) that carries *3x-FLAG-fzlA* as the sole copy at the native locus and subjected the eluate to mass spectrometry (MS)-based protein identification, using WT cells (not producing Flag-tagged FzlA) as a negative control. MS identified FtsZ, a known interactor for FzlA, in the 3xFLAG-FzlA eluate, validating this approach (Table S2). From the proteins identified and enriched more than 5-fold in the 3xFLAG-FzlA eluate, we identified FtsK, a known cell division protein, as a putative binding partner (Table S2).

FtsK is an essential, bifunctional cell division protein with an N-terminal domain (FtsK_N_: residues 1-213) containing four transmembrane passes and a cytoplasmic C-terminal domain (FtsK_C_: residues 315-825), connected by a disordered linker (residues 213-315) (Wang et al., 2006). FtsK_N_ is responsible for localizing FtsK to the divisome and plays an essential role in constriction, while FtsK_C_ is a hexameric DNA translocase that segregates the chromosomal termini into each daughter cell during division. To confirm our MS results, we generated a strain in which *mChy-ftsK* was expressed at the *ftsK* locus as the sole copy, in a WT (EG2427) and *3xFLAG-fzlA* background (EG2428) and performed anti-FLAG coIP followed by immunoblotting. Western blot analysis revealed that 3xFLAG-FzlA interacted with mChy-FtsK *in vivo* (Fig. 7A). To pursue the interaction further, we performed reciprocal coIP in a strain which harbored *FLAG-ftsK* at the *ftsK* locus as the sole copy (EG742) and compared to WT. Western blot analysis demonstrated that FLAG-FtsK interacts with native, untagged FzlA *in vivo* as well (Fig. 7B).

**Figure 7.**
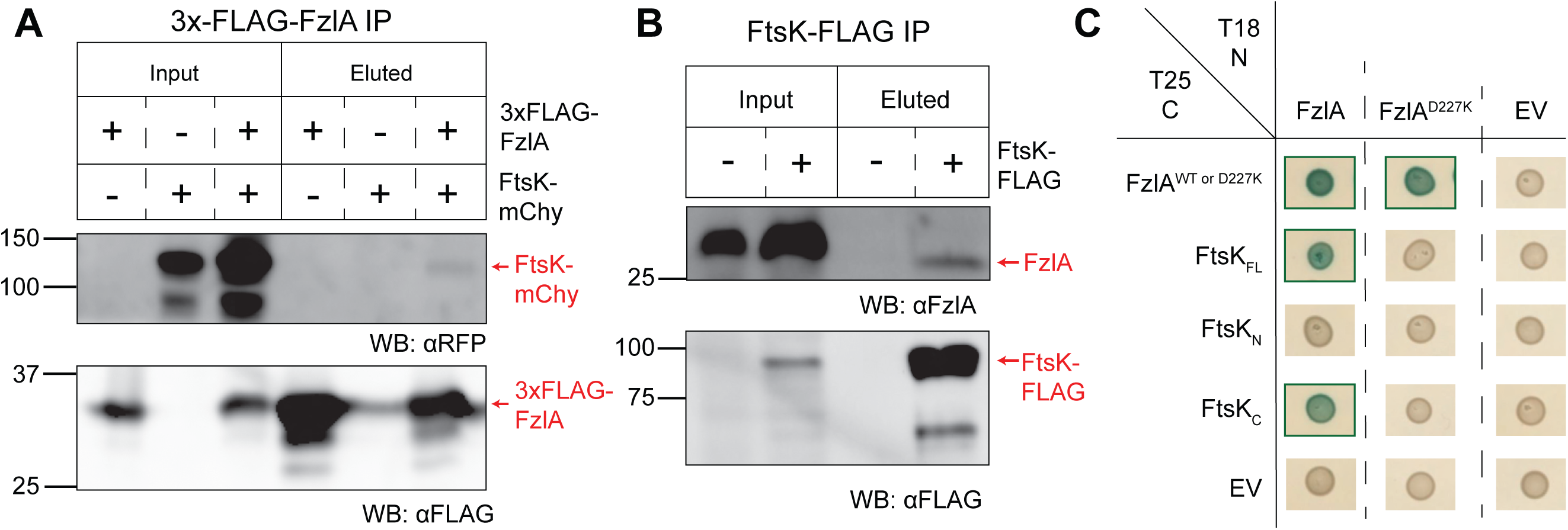
FzlA interacts with FtsK_C_ via the FzlA C-terminal tail. **A-B.** Western blots of whole cell lysates or eluates from an immunoprecipitation (IP) using α-FLAG resin. Ladder values to the left of the blots are in kDa. Red arrows to the right of the blots indicate to what protein the band is attributed. Presence or absence of the fusion proteins in the strains used are indicated above the blots by plus (+) or minus (-) signs, respectively. **A.** 3xFLAG-FzlA immunoprecipitation. Blots were incubated with primary antibodies recognizing either mCherry (αRFP) or FLAG (αFLAG). Lane 1 and 4, *fzlA*::*3xFLAG-fzlA* (EG2217). Lane 2 and 5, *ftsK*::*mCherry-ftsK* (EG2427). Lane 3 and 6, *fzlA*::*3xFLAG-fzlA*; *ftsK*::*mCherry-ftsK* (EG2428). **B.** FtsK-FLAG immunoprecipitation. Blots were incubated with primary antibodies recognizing either FzlA (αFzlA) or FLAG (αFLAG). Lane 1 and 3, WT (EG865). Lane 2 and 4, *ftsK*::*ftsK-FLAG* (EG742) **C.** Bacterial Two-Hybrid results for interaction between FzlA variants and full-length FtsK or its domains. The adenylyl cyclase subunits T18 (left) and T25 (top) are fused to proteins via the N-terminus or C-terminus, respectively. A green box around the representative spot image means that the three biological triplicates were positive for induction of the cAMP-dependent β-galactosidase reporter, indicating a positive interaction. EV, empty vector. FtsK_FL_, Full-length FtsK. FtsK_N_, FtsK N-domain. FtsK_C_, FtsK C-domain.

### The FzlA-FtsK interaction occurs between the FzlA C-terminal tail and FtsK_C_

To narrow down the domains responsible for the FzlA-FtsK interaction, we performed Bacterial Two-Hybrid (BTH) analysis. We first sought to confirm our coIP-MS results for full-length, WT FzlA and FtsK using BTH. FzlA forms a homodimer and exhibits self-interaction by BTH, allowing FzlA self-interaction to serve as an internal positive control (Fig. 7C). Indeed, FzlA and full-length FtsK (FtsK_FL_) were able to interact by BTH, suggesting a direct interaction in this heterologous system (Fig. 7C).

In narrowing down the interacting surfaces on FzlA, we sought to test residues previously determined to be essential for FzlA function. We hypothesized that if the FzlA-FtsK interaction is essential, the downstream signaling would likely occur through the essential C-terminal tail (Lariviere et al., 2018). To test this hypothesis, we used a point mutant in the penultimate residue of FzlA (FzlA^D227K^), a variant that is unable to function in division despite retaining its ability to bind to and colocalize with FtsZ (Lariviere et al., 2018, 2019). Consistent with our hypothesis, FzlA^D227K^ was unable to interact with FtsK_FL_ by BTH, but was still able to self-interact, suggesting the protein is properly folded but that the C-terminus is required to bind FtsK. (Fig. 7C).

FtsK_N_ and FtsK_C_ have independent, separable, essential functions *in vivo* (Wang et al., 2006). To determine which domain of FtsK interacts with FzlA, we tested each separately for its ability to interact with FzlA by BTH. The FtsK_N_ construct included the disordered linker up to residue 333, while FtsK_C_ included the remainder of the protein. FzlA was able to interact with FtsK_C_, but not with FtsK_N_ (Fig. 7C). To determine if the FzlA C-terminal tail is required for the interaction, we tested the ability of FtsK_C_ and FzlA^D227K^ to interact. FzlA^D227K^ and FtsK_C_ were able to interact by only one orientation, suggesting reduced affinity when the C-terminal tail is mutated (Fig. 7C, Fig. S5). (Fig. S4). Collectively our coIP-MS and BTH approaches indicate that FzlA and FtsK directly interact via the C-terminal tail of FzlA and the DNA translocase domain of FtsK.

### FtsK functions upstream of FtsWI

Prior work characterizing *C. crescentus* FtsK demonstrated that depleting the C-terminal domain of FtsK causes cell chaining, with multiple cell bodies connected by unresolved constrictions, whereas depleting full-length FtsK causes cell filamentation without constriction (Wang et al., 2006). We hypothesized that if FtsK acts downstream of FzlA in an FtsWI activation pathway, in the *ftsW**I\** background where *fzlA* is non-essential FtsWI should also be less dependent on FtsK-mediated activation. While *ftsK* remains essential in a *ftsW**I\** background based on comparative transposon sequencing (Lariviere et al., 2019), we reasoned that examining the phenotype associated with depletion of FtsK in a *ftsW**I\** background could reveal a genetic relationship between *ftsK* and *ftsWI*. We generated strains with xylose-dependent expression of *ftsK* in which *ftsK_FL_* was placed at the *xylX* locus and native *ftsK* was replaced with a spectinomycin resistance cassette at the native locus, in both the WT (EG3332) and *ftsW**I\** (EG3333) backgrounds. We also examined the phenotype of the previously characterized FtsK_C_ depletion strain (LS4202/EG2996) (Wang et al., 2006).

We confirmed the previously observed depletion phenotypes for FtsK_FL_ and FtsK_C_, which were predominantly filamentous and chained cells, respectively (Fig. 8A, chaining: blue arrows). In liquid media, cells depleted of full-length FtsK in a WT background still experienced an increase in absorbance, likely owing to continued growth of filamentous cells. However, FtsK depletion was lethal on solid media (Fig. 8, B-C). We reasoned that if FtsK plays a role downstream of FzlA in activating FtsWI, we might observe constrictions or even completed division events in cells depleted of FtsK in the *ftsW**I\** background. This expectation would be similar to our observations that FzlA depletion in a WT background leads to filamentation, whereas *ftsW**I\** cells lacking FzlA still divide (Lariviere et. al, 2019). Indeed, cells depleted of FtsK_FL_ in a *ftsW**I\** background were not filamentous. Instead, they were short, with pleiotropic phenotypes (Fig. 8A) including thin, extended constrictions (magenta arrows); chaining (blue arrows); and/or lysis (orange arrows). The phenotypes of cells depleted of FtsK_FL_ in the *ftsW**I\** background were reminiscent of those associated with long-term *fzlA* over-expression in the *ftsW**I\** background. Despite the presence of cells of normal length, FtsK depletion in the *ftsW**I\** background was still lethal, indicating an essential function of FtsK in addition to its role in regulating the activation state of FtsWI. These results collectively indicate that FtsK lies between FzlA and FtsWI in a constriction activation pathway and that regulatory input from FtsK may couple chromosome segregation to constriction activation.

**Figure 8.**
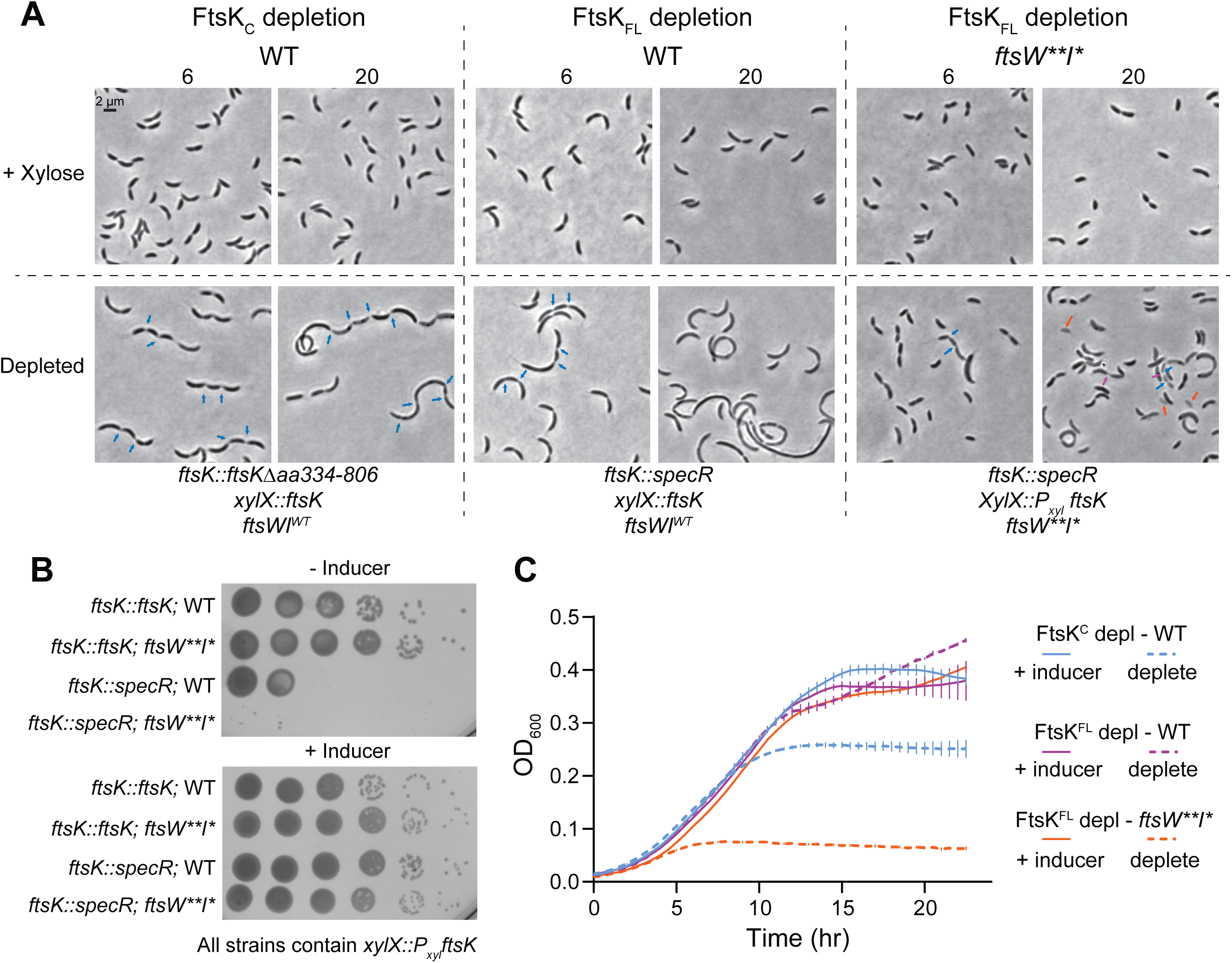
FtsK acts upstream of FtsWI and is required for constriction initiation. **A.** Representative phase contrast of strains during depletion of the FtsK C-domain (FtsK_C_) or full-length FtsK (FtsK_FL_) in either wild-type or *ftsW**I\** backgrounds. Cells were grown in the presence of 0.3% xylose (top row, *ftsK* induced) or 0.2% glucose (bottom row, FtsK or FtsK_C_ depleted) with images taken at six and 20 hours post depletion. FtsK_C_ depletion, EG2996. FtsK_FL_ depletion in WT, EG3332. FtsK_FL_ depletion in *ftsW**I\**, EG3333. **B.** Spot dilutions of strains with xylose-inducible *ftsK* expression in the wild-type or *ftsW**I\** background, in the absence or presence of native *ftsK*. Plates contain either 0.2% glucose (- Inducer) or 0.3% xylose (+ Inducer). Row 1, EG3299. Row 2, EG3300. Row 3, EG3332. Row 4, EG3333. **C.** Growth curves of strains dependent on xylose for FtsK expression (same as A). Solid lines indicate grown in the presence of 0.3% xylose. Dashed lines indicate grown in the presence of 0.2% glucose. FtsK_C_ depletion, EG2996. FtsK_FL_ depletion in WT, EG3332. FtsK_FL_ depletion in *ftsW**I\**, EG3333. Points are the mean of three technical replicates at that timepoint, error bars are the standard error of the mean. Shown is a representative replicate of a biological triplicate.

### Proper regulation of FzlA-mediated signaling is critical to prevent DNA damage

Cell chaining is observed in strains that have either cell separation (Meier et al., 2017) or chromosome segregation defects (Wang et al., 2006; Yu et al., 1998). Identification of a FzlA-FtsK interaction led us to hypothesize that the cell chaining observed during both FtsK depletion and *fzlA* overexpression in the *ftsW**I\** background is due to defects in coupling chromosome segregation to division. Specifically, we hypothesized that misregulation of FzlA-FtsK-mediated signaling in a background with hyperactive constriction traps DNA at the division site, preventing the completion of division and causing DNA damage. To test this, we leveraged a previously characterized *P_sidA_-egfp* reporter construct (Modell et al., 2014). The *sidA* promoter is sensitive to DNA damage, leading to production of EGFP when DNA damage occurs. To obtain a baseline of DNA damage in *Caulobacter*, we introduced the reporter construct into a WT background (EG3653). Since DNA damage is rare in WT cells without stress, expression was relatively low. To compare among different strains, we set a threshold for “high GFP” as above the 99^th^ percentile of WT GFP values. Any cell with a value higher than that threshold was considered to have high GFP and, by proxy, DNA damage.

To understand how hyperconstriction and the presence of FzlA influence DNA damage, we introduced the DNA damage reporter construct into *ftsW**I\** (EG3655) and *ΔfzlA, ftsW**I\** (EG3667) backgrounds. Consistent with our hypothesis, the hyperconstricting *ftsW**I\** population had a modest increase in DNA damage (Table S3, Fig. 9A, 4.59%) (Modell et al., 2011, 2014). Interestingly, the *ΔfzlA, ftsW**I\** population had WT levels of DNA damage (Table S3, Fig. 9A, 0.79%), which suggests that hyperconstriction is the underlying cause of DNA damage (since *ΔfzlA* cells constrict more slowly than WT or *ftsW**I\**), not insensitivity of FtsW**I* to the DNA damage inhibitors SidA and DidA (Modell et al., 2014, 2011).

**Figure 9.**
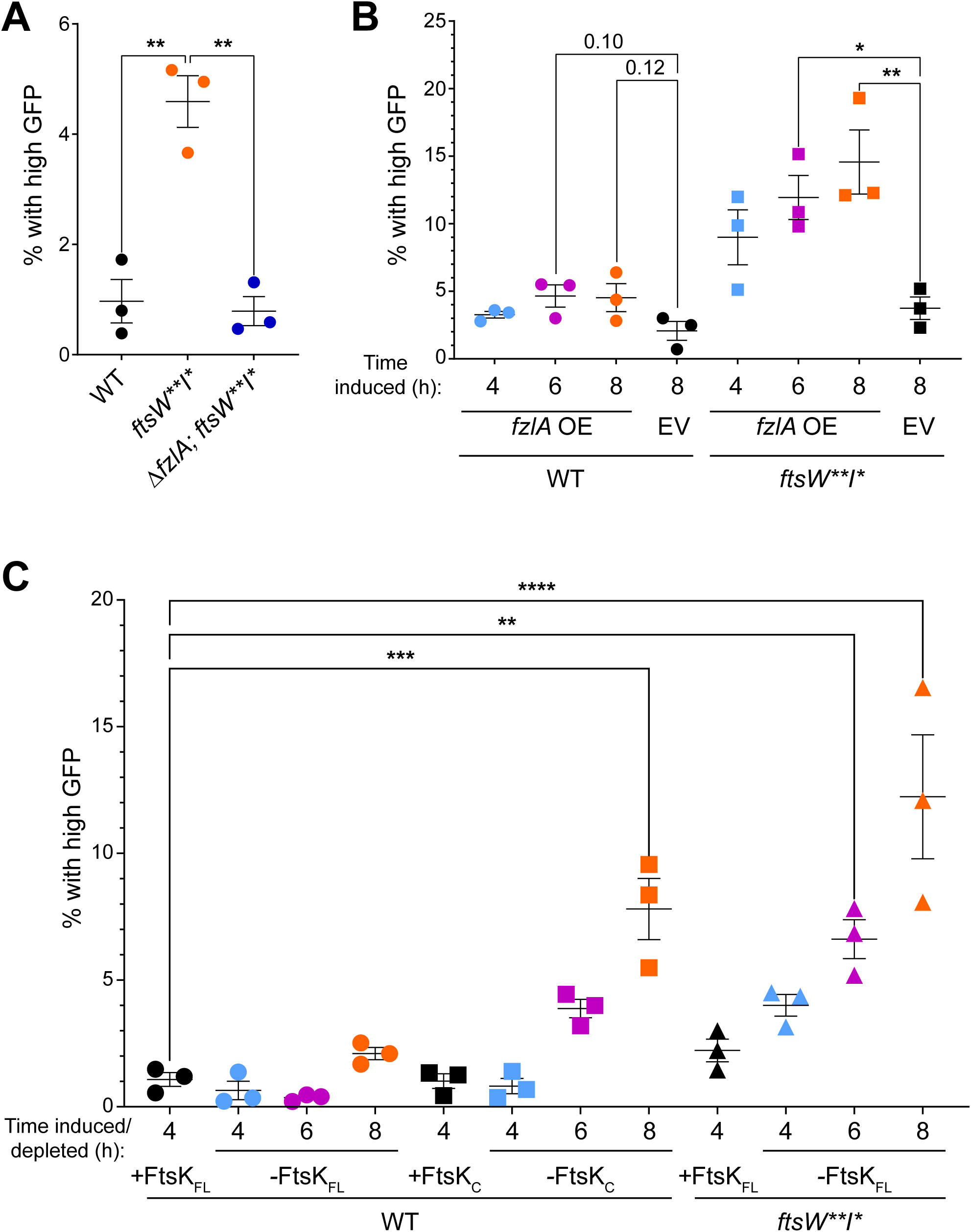
Misregulation of the FzlA-FtsK-FtsWI pathway results in DNA damage. **A-C.** Plot of the percentage of cells containing high levels of EGFP (greater than the 99^th^ percentile of the specified control strain) from the *P_sidA_-egfp* reporter. Plots comparing high EGFP producing cells between **A.** wild-type (WT, EG3653), *ftsW**I\** (EG3655), and *ΔfzlA*; *ftsW**I\** (EG3667) backgrounds or **B.** strains containing either an empty vector (EV) control or *fzlA* overexpression (OE) construct in WT (EV: EG3658, *fzlA* OE: EG3664) or *ftsW**I\** (EV: EG3666, *fzlA* OE: EG3672) backgrounds in the presence of 0.3% xylose for four, six, and eight hours. * represents a p-value < 0.05. Averages of biological triplicates are shown and were used for statistical analysis. **C.** Plot comparing high EGFP-producing cells grown in 0.3% xylose for 4 hr of a full-length FtsK_FL_ depletion strain in a WT background (EG3904 + xyl 4hr) to various depletion strains grown in the presence or absence of 0.3% xylose (EG3904: FtsK depletion in WT, EG3900: FtsK_C_ depletion in WT, EG3902: FtsK depletion in *ftsW**I\**) at four, six, and eight hours. * represents a p-value < 0.05. Averages of biological triplicates are shown and were used for statistical analysis.

To determine if hyperconstriction due to an overabundance of FzlA also leads to DNA damage, we introduced the DNA damage reporter into our xylose-dependent *fzlA* over-expressing and EV strains in the WT and *ftsW**I\** backgrounds. The EV control in a WT (EG3658) background had WT levels of DNA damage when supplemented with 0.3% xylose for eight hours (Table S3, Fig. 9B, 2.1%). The FzlA-overproducing WT strain (EG3664) experienced a modest increase in the amount of EGFP expression, starting at 3.3% at four hours of over-production and increasing to near 4.6% at six and eight hours (Table S3, Fig. 9B). The maximum levels of DNA damage we observed for FzlA overproduction were similar to the levels observed in the *ftsW**I\** background.

We next assessed DNA damage in cells overproducing FzlA (EG3672) compared to the EV control (EG3666) in the *ftsW**I\** background. Considering that this is the background in which FzlA overproduction led to cell chaining, we hypothesized the DNA damage may be much greater than in the WT background. In line with this hypothesis, four hours of FzlA over-production in the *ftsW**I\** background significantly increased DNA damage when compared to the EV control (Table S3, Fig 9B, 9 vs 4.6%). The amount of DNA damage increased with longer induction times (Table S3, Fig 9B, 11.94% at 6 hr and 14.56% at 8 hr), consistent with our observations of cell chaining at six hours post-induction. We conclude that hyperactivation of constriction by FzlA overproduction leads to DNA damage.

We next tested whether misregulating cell division through modulation of FtsK, itself, causes DNA damage. To obtain a baseline of DNA damage for this set of experiments, we introduced the DNA damage reporter construct into the xylose-dependent FtsK_FL_ depletion strain (EG3332) described above (Fig. 8). We set a threshold for “high GFP” as above the 99^th^ percentile of GFP values observed in this strain grown in the presence of xylose to induce *ftsK_FL_* expression. We then introduced the DNA damage reporter construct into the FtsK_FL_ (EG3904) and FtsK_C_ (EG3900) depletion strains in a WT background. We hypothesized that depleting FtsK_FL_ in a WT background would not result in DNA damage since these cells fail to initiate constriction and become filamentous. Supporting our hypothesis, we observed little to no increase in DNA damage at 4 (Table S3, Fig. 9C, 0.647%), 6 (Table S3, Fig. 9C, 0.360%) and 8 hours of depletion (Table S3, Fig. 9C, 2.10%) when compared to the replete condition (Table S3, Fig. 9C, 1.08%). We next examined the effect of depleting FtsK_C_ as this results in cell chaining. Indeed, there were higher levels of DNA damage compared to our replete control strain (Table S3, Fig. 9C, 1.08%) at 6 (Table S3, Fig. 9C, 3.88%) and 8 hours (Table S3, Fig. 9C, 7.80%) of depletion, with DNA damage increasing over time as FtsK_C_ was depleted.

Finally, we tested the effect of depleting FtsK_FL_ in an *ftsW**I\** background since these cells experience chaining and extended constrictions. We introduced the DNA damage reporter construct into a *ftsW**I\** strain with the only copy of *ftsK_FL_* under the control of the xylose promoter (EG3902). In contrast to FtsK depletion in a WT background, when we depleted FtsK in this background we observed an increase in DNA damage compared to our control strain (Table S3, Fig. 9C, 1.08%) at 4 (Table S3, Fig. 9C, 4.00%) 6 (Table S3, Fig. 9C, 6.62%), and 8 hours of depletion (Table S3, Fig. 9C, 12.2%). Collectively, these data indicate that misregulating the FzlA-FtsK-FtsWI pathway results in DNA damage due to improper coupling of chromosome segregation to constriction, and reinforces our hypothesis that FtsK plays a central role in this constriction activation process.

## Discussion

The specifics of bacterial constriction activation can vary from species to species, but the functional requirements to select the division plane, dynamically position the division components, segregate the chromosomes, and activate constriction in a controlled manner are essential and conserved. Our work here describes a key role for FzlA in signaling to activate FtsWI in coordination with chromosome segregation. FzlA activates constriction during division by converting fast-moving FtsW molecules into an active, slow-moving state (Figs. 1-5). The FzlA activation pathway signals through FtsK, an essential protein that segregates sister chromosomes during division (Figs. 7-8). When this signaling is misregulated through hyperactivating mutations or over-production of FzlA, the cell suffers from cell envelope defects (Lariviere et al., 2019; Modell et al., 2014) (Fig. 6) and DNA damage (Fig. 9), likely due to defects in coordinating chromosome segregation with division. We propose that, prior to activation, inactive FtsWI is either stationary or is fast-moving through indirect association with treadmilling FtsZ clusters (Fig. 10A). FzlA binds FtsZ and may infrequently move by dynamic association with FtsZ filament ends or, more frequently, remain stationary (but treadmilling) by association with the interior of FtsZ filaments. Upon binding the C-domain of FtsK through its C-terminal tail, FzlA signals through FtsK to license FtsWI activation, likely acting through additional factors. Ultimately FtsWI is converted from a fast-moving, inactivate state to an active state that moves slowly, guided by PG synthesis (Fig. 10B). FzlA can move together with the activated FtsWI complex.

**Figure 10.**
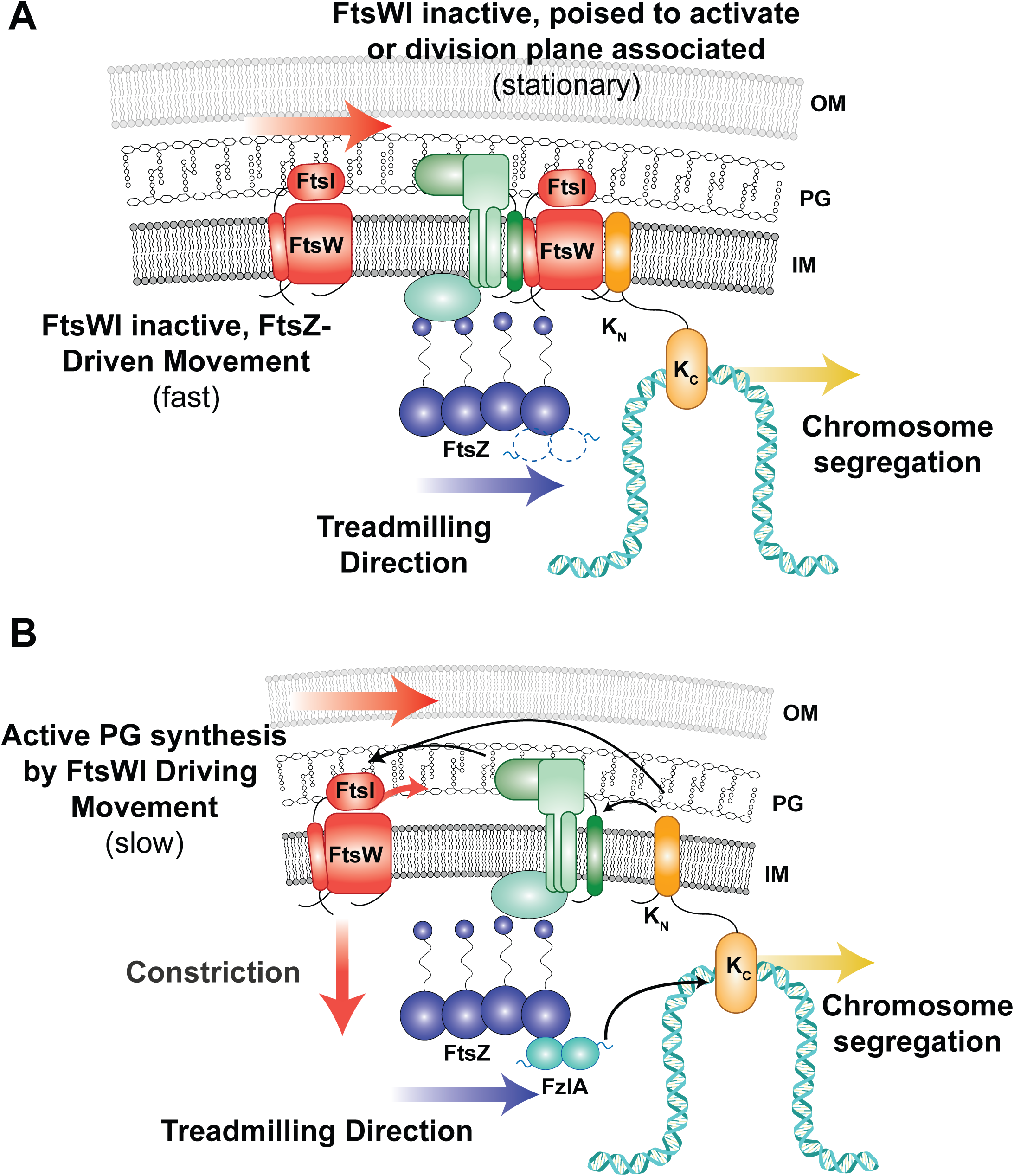
FzlA binds to FtsZ and signals through FtsK to shift inactive FtsW to a slow-moving, active state. **A.** Inactive FtsW molecules either move with FtsZ clusters as they treadmill about the division plane or are stationary, either because they are poised to activate or bound to a stationary partner at the division plane. FtsK begins segregating chromosomal termini after localizing. **B.** FzlA binds to FtsZ through the FtsZ-binding residues while the FzlA C-terminal tail binds to the FtsK_C_ to signal downstream for FtsWI activation, after properly ensuring the division plane is free of obstruction by DNA.

Our findings provide a deeper understanding of the mechanisms of FtsWI hyperactivation and its consequences. We observed that active FtsW** synthesized PG for longer periods than FtsW. Conversely, FzlA did not impact the processivity of active FtsW or FtsW**, suggesting increased processivity results from a mechanism distinct from that of FzlA-mediated activation. The mutations of FtsW** may perturb association with a regulator that modulates processivity. An obvious candidate is the FtsQLB complex, based on the observations made in *E. coli* and *P. aeruginosa* (Tsang and Bernhardt, 2015; Marmont and Bernhardt, 2020; Park et al., 2020), though further experiments are required to test this hypothesis. Alternatively, mutation of FtsWI may directly impact processivity of the enzyme.

Our study also illuminates characteristics of stationary PG synthases. In the presence of FzlA, stationary FtsW** remain stationary for longer times than FtsW. Because FzlA is present and FtsW** has a greater propensity for the active state, we do not believe that these long-lived molecules are poised for activation. One hypothesis is that FtsW** has increased affinity for the cell wall, which is not moving. It is also possible that FtsW** has increased affinity for a divisome component that remains stationary. In *B. subtilis,* the FtsZ-binding proteins have been observed to be largely stationary (Squyres et al., 2021), so a candidate for this role is one of the FtsZ membrane anchors. The proportion of stationary FtsW or FtsW** was also largely unaffected by depletion or overproduction of FzlA, except in the case of FtsW** in the absence of FzlA. Since FtsW** is primed for activation, we hypothesize that being poised for activation, but awaiting FzlA for a prolonged period results in inactive, moving FtsW shifting to a stationary state. Moreover, in the absence of FzlA, both FtsW (FzlA depletion) and FtsW** (*ΔfzlA*) have shorter stationary times than when FzlA is present, suggesting that short-lived stationary molecules may be those that are poised to be activated. Collectively, our results suggest one stationary population of FtsW is short-lived and awaiting activation by FzlA and another population has longer-lived associations with a stationary fixture at the division plane.

We discovered that FzlA converts inactive, fast-moving FtsW to an active, slow-moving state and that upstream signaling from FzlA is limiting for FtsW activation. Our observations for FzlA are, in some ways, similar to observations made for FtsN in hyperactive *ftsB* mutant (*ftsB^E56A^)* and WT backgrounds in *E. coli* (Yang et al., 2021). Expression of *ftsB^E56A^*hyperactivates the PG synthases, causes cell shortening, and renders *ftsN* non-essential. The double mutant of *ΔftsN, ftsB^E56A^*, results in cell lengths more similar to WT *E. coli* and the PG synthases are active less often than in the *ftsB^E56A^* background with FtsN. FtsN depletion in an otherwise WT background also results in cell filamentation and the PG synthases are more often inactive, consistent with FtsN being an FtsW activator. These results mirror many of our FzlA results and may be a common feature of factors that serve to activate FtsWI. However, there are notable differences between FzlA and FtsN and we propose that they act at distinct stages of divisome activation in α-proteobacteria to couple division to other cellular processes. FtsN is transmembrane protein with a PG-sensing SPOR domain that is proposed to couple PG synthesis to amidase activity at the division site (Yahashiri et al., 2015; Gerding et al., 2009). Based on genetics in *E. coli*, FtsN is thought to bind directly to the FtsQLB complex which directly interacts with and activates FtsWI (Park et al., 2021; Gerding et al., 2009; Marmont and Bernhardt, 2020). Conversely, FzlA is a soluble FtsZ-binding protein that also binds the chromosome-segregation factor FtsK. Considering that *C. crescentus* has essential homologs of FtsN and FtsQLB, we propose that FzlA and FtsK provide an additional layer of regulation to FtsN-FtsQLB-mediated activation that serves to integrate information from the Z-ring and chromosome with constriction.

In *C. crescentus,* the essential divisome components include FtsZ, FtsA, FzlA, FtsK, FtsN, FtsQLB, and FtsWI (Goley et al., 2011). FzlA is restricted to the α-proteobacteria and FtsN is restricted to proteobacteria, but the remainder of these proteins are widely conserved across bacterial phyla. Our work characterizing the FzlA-FtsK interaction identifies, for the first time, where FtsK integrates in the constriction activation pathway in α-proteobacteria. The FzlA-FtsK interaction is likely conserved throughout α-proteobacteria, as residues 223, 227, and 228 of the FzlA C-terminal tail are invariant in FzlA homologs (Lariviere et al., 2018, 2019). FzlA homologs are even found in the Rickettsiales, which have streamlined genomes and are often outliers among the α-proteobacteria in conservation of factors that are essential in *C. crescentus*. Outside of α-proteobacteria, FtsK and the family of FtsK/SpoIIIE/HerA homologs are widespread and often essential, even extending to archaea (Bigot et al., 2007). In *E. coli,* only FtsK_N_ is essential, which suggests that its role in the constriction activation pathway is conserved, but that redundant chromosome maintenance mechanisms support growth in the absence of FtsK_C_ (Draper et al., 1998). Notably, *E. coli* encodes a nucleoid occlusion system as an additional chromosome maintenance mechanism. SlmA, the protein responsible for nucleoid occlusion in *E. coli*, binds to the nucleoid and prevents proper Z-ring assembly wherever it is present, which prevents the cell from constricting onto the nucleoid (Bernhardt and de Boer, 2005). *C. crescentus* and most other α-proteobacteria lack nucleoid occlusion systems. We therefore propose that FzlA functions in this group to robustly couple constriction to chromosome segregation through FtsK.

Our findings highlight distinct mechanisms of chromosome maintenance that ultimately regulate constriction in different bacteria. Our work suggests that, in α -proteobacteria, the FtsZ-FzlA-FtsK pathway coordinates constriction with chromosome dimer resolution and segregation of termini, while other bacteria rely more heavily on negative regulators of FtsZ at the earliest stages of Z-ring assembly. Once chromosomes are segregated, the FzlA-FtsK interaction promotes a shift of FtsWI to an active state. FzlA, and presumably FtsK (likely when it is not associated with the chromosome), travel with FtsW for a time, but there are likely additional mechanisms influencing FtsWI processivity. Future experiments will identify how other components contribute to regulation of constriction activation and will further clarify the mechanistic underpinnings of the FtsZ-FzlA-FtsK activation pathway to FtsWI.

## Methods

### *C. crescentus* and *E. coli* growth media and conditions

*C. crescentus* NA1000 cells were grown at 30°C in peptone-yeast extract (PYE) media. *E. coli* NEB Turbo (NEB Catalog #C2986K) and BTH101 cells were grown at 37°C in Luria-Bertani (LB) medium. Xylose was used at final concentrations of either 0.001% or 0.3% (w/v) as indicated. Glucose was used at final concentrations of either 0.001% or 0.2% (w/v). For depletion strains, cells were grown in PYE supplemented with xylose as indicated prior to being washed with plain PYE medium three times and resuspended in the PYE medium supplemented with the stated concentrations of glucose or xylose. Solid media included 1.5% (w/v) of agar. Antibiotics were used in liquid (solid) medium at the following concentrations for *C. crescentus* growth: kanamycin, 5 (25) µg/mL; gentamycin, 1 (5) µg/mL; spectinomycin, 25 (100) µg/mL; and oxytetracycline, 1 (2) µg/mL. Streptomycin was used at 5 µg/mL in solid media. *E. coli* antibiotics were used in liquid (solid) medium as follows: kanamycin, 30 (50) µg/mL; gentamycin 15 (20) µg/mL; spectinomycin, 50 (50) µg/mL; ampicillin, 50 (100) µg/mL; and oxytetracycline, 12 (12) µg/mL. For growth curves, a Tecan Infinite M200 Pro plate reader measured absorbance every 30 minutes at OD_600_ of a 100 µL culture volume, at a starting OD_600_ of 0.05, in a 96-well plate in biological and technical triplicate with intermittent shaking. In the growth curve of Figure 2.3 C, cultures were pre-induced for 6 hours before spotting. For spot dilution assays, mid-log cells were diluted to an OD_600_ and serially diluted up to 10^-5^ before spotting 5 µL onto a PYE plate with indicated inducer and/or antibiotic. Plates were incubated at 30°C for 48 hours. For time-lapse microscopy or single-molecule tracking experiments, induction with 0.3% xylose occurred for one hour before synchrony and throughout the length of the experiment.

### Synchrony

Synchrony was performed as previously described (Goley et al., 2011). Cells were grown overnight in 15 mL of PYE medium to an OD_600_ between 0.1 and 0.5, harvested by pelleting at 6,000 rpm at 4°C, resuspended in 750 µL of ice-cold M2 salts, and combined with 750 µL of Percoll (Fisher Scientific Catalog #45-001-748). The cell suspension was centrifuged at 20,000 x g for 20 minutes at 4°C. The swarmer (lower) band was isolated, transferred to a fresh tube, and pelleted at 8,000 x g for 2 minutes. The swarmer pellet was washed twice with 1 mL of ice-cold M2 salts, and pelleted each time at 8,000 x g for 2 minutes. The swarmer cells were resuspended in PYE with appropriate additives and grown at 25°C. Samples were analyzed by phase-contrast or fluorescence microscopy.

### Phase-contrast and standard epifluorescence microscopy

Cells in exponential phase of growth were spotted on pads made of 1% agarose resuspended in water or PYE medium, supplemented with 0.3% xylose when appropriate, and imaged using a Nikon Eclipse Ti inverted microscope equipped with a Nikon Plan Fluor 100X (NA1.30) oil Ph3 objective and Photometrics CoolSNAP HQ^2^ cooled CCD camera. Timelapse imaging was performed as previously described (Lariviere et al., 2019), using Gene Frames (Thermo Fisher Scientific Catalog #AB0577) to ensure a tight seal during imaging. After synchrony, cells were resuspended in PYE with the appropriate additives and a stopwatch was started to begin measuring pre-constriction time. The cell suspension was spotted on agarose pads. After the pad was allowed to dry sufficiently, the top coverslip was adhered to the gene frame and timelapse imaging was initiated. The timelapse imaging proceeded by taking an image every five minutes for 4 hours. Images were processed using FIJI with the MicrobeJ plugin or using Adobe Photoshop.

### Timelapse microscopy analysis for growth metrics comparison

To determine dimensions of log-phase cells, cell length and width were measured using FIJI (Schindelin et al., 2012) and MicrobeJ software, similar to as previously described (Lariviere et al., 2018). Constriction rate and elongation rate were also determined using MicrobeJ (Ducret et al., 2016). MicrobeJ software allowed for tracking of cells imaged by time-lapse microscopy throughout the division process, with automatic detection of constriction initiation and manual determination of cell separation. Cell length was found for cells at each time point, cell width was found at the site of constriction, and constriction time was calculated by multiplying the number of frames in which constriction was detected by 5 (since images were acquired every 5 minutes), allowing for calculation of constriction and elongation rates. Lethal division events were quantified manually by counting dividing cells that had 1) one daughter halt growth, 2) both daughters halt growth, and 3) one daughter lyse post-division. Prism was used for graphing and statistical analysis of calculated terms. Statistical analysis for Superplots involved one-tailed Mann-Whitney tests, based on the hypothesis that FzlA overproduction would enhance constriction. All cells that were in focus were included in these analyses.

### Immunoblot analysis

Immunoblot analysis was performed using standard procedures, with a 1:6,666 dilution of affinity-purified α-FzlA antibodies (Goley et al. 2010), a 1:2,500 dilution of α-MreB antisera (Beaufay et al 2015), a 1:2500 dilution of α-RFP antisera (Chen et al. 2005), a 1:1000 dilution of α-FLAG M2 monoclonal antibodies and/or 1:10,000 dilution of HRP-labeled α-rabbit secondary antibodies (BioRAD Catalog #170-6515) on nitrocellulose membranes. Clarity Western Electrochemiluminescent substrate (BioRAD Catalog #170-5060) was added to facilitate protein visualization via an Amersham Imager 600 RGB gel and membrane imager (GE).

### Advanced epifluorescence imaging and single-molecule tracking of Halo-FtsW and Halo-FzlA

Except for the FzlA depletion experiment, all single-molecule tracking (SMT) experiments were performed after synchrony. After synchrony, cells were labeled with Janelia Fluor 646 (a gift from Dr. Luke Lavis, Howard Hughes Medical Institute) at an empirically determined concentration for each sample to obtain single-molecule level labeling, ranging from 0.2 nM to 10 nM in PYE with additives as required (xylose or glucose). The cell suspension was incubated in the dark with shaking at 25°C for 15 minutes before washing away the dye with three washes of 1 mL PYE. Cells were resuspended in fresh PYE and allowed to continue growing until the time of an experiment. Cells were imaged no later than 90 minutes after synchrony, with the imaging beginning one-hour post-synchrony.

SMT experiments were performed as previously described (Yang et al., 2021). Briefly, single JF646-labeled Halo-FtsW or Halo-FzlA molecules were detected on an Olympus IX-81 inverted microscope equipped with an UPlanApo 100x TIRF (1.50NA/oil) objective and an Andor iXon 897 Ultra EM-CCD camera in epifluorescence-illumination mode using a Coherent 647-nm laser at ∼2 W cm^-2^ power density. Brightfield and fluorescent images were acquired using Metamorph 7.8.13.0 software. The Camera’s emGain was turned to 300 with a pre-amplifier gain setting on 3 and digitizer set to 16-bit. Baseline clamp was activated, with a baseline offset set to 100. The focal plane was placed at the middle of the cell to capture the most moving molecules at the division plane. SMT images were taken continuously with 500-ms exposure time for 200 s.

Fluorescent spots of single FtsW or FzlA molecules were first localized using two-dimensional Gaussian fitting in an ImageJ (1.52p) (Schneider et al., 2012) plug-in, ThunderSTORM (dev-2016-09-10-b1) (Ovesný et al., 2014). A bandpass filter (80–250 nm) for sigma was applied to remove the single-pixel noise (small sigma) and out-of-focus (big sigma) molecules. To focus on the FtsW or FzlA involved during cell division, especially cytokinetic PG synthesis, we only analyzed the molecules located at mid-cell or the places with observable invagination. In the case of the FzlA depletion strain (EG3523) without obvious constrictions, ZapA-mNeonGreen-marked Z-rings were used to identify divisome-associated FtsW trajectories. Those localizations were then linked to trajectories using a home-built Matlab 2020a script (https://github.com/XiaoLabJHU/SMT_Unwrapping) adopting the nearest neighbor algorithm as previously described (Sbalzarini & Koumoutsakos, 2005) https://www.nature.com/articles/s41564-020-00853-0 - ref-CR49. For the movement of each molecule, the rate for each segment was measured and recorded, but for calculating the proportion of moving molecules, the molecule was counted once.

The unwrapped and corrected trajectories described above were then segmented to determine whether Halo-FtsW or Halo-FzlA molecules in a segment are stationary or moving directly as previously described (Yang et al., 2021). The cumulative probability density function of the directional moving speeds was further calculated and fit to a two-population model (of log-normal distribution) as described previously (Yang et al., 2021). In this work, we did not fix the fast-moving speed since the FtsZ treadmilling speed is undetermined in *Caulobacter*.

### Co-immunoprecipitation

Co-immunoprecipitation was performed as previously described (Bowman et al., 2008; Lariviere et al., 2018). All coIP steps, unless designated, were performed at 4°C. CoIP buffer consists of 20 mM HEPES pH 7.5, 100 mM NaCl, and 20% glycerol and one Pierce Protease Inhibitor Mini Tablet, EDTA Free, (Fisher Scientific, A32955) per liter of coIP buffer. Cells were harvested during exponential growth by centrifugation at 6,000 x g at 4°C. Cells were washed with coIP buffer, pelleted at 6,000 x g, and resuspended in coIP buffer. Dithiobis(succinimidyl propionate) (DSP) was used to crosslink free amines. DSP was resuspended at 10 mg/mL in DMSO and added to the cell suspension to reach a final concentration of 2 mM of DSP in 0.08% DMSO for 1 hr at 4°C and 10 min at room temperature. The reaction was quenched by adding 1 M Tris-HCl, pH 7.5 to a final concentration of 20 mM for 10 minutes. After quenching, cells were lysed by three passes an EmulsiFlex-C3 at 16,000 psi. Cell membranes were solubilized by adding sodium deoxycholate to 0.5% w/v, Igepal CA-630 to 1% w/v, and EDTA to a 2 mM final concentration. Insoluble cellular debris was removed by pelleting at 4,500 x g for 10 min. For each sample, up to 40 mL of supernatant of each sample was transferred to a fresh tube and normalized based on OD to the lowest density sample. To each tube, 100 µL of washed α-FLAG resin (Sigma-Aldrich, A2220-5ML) was added before incubating on a rotator overnight. Afterwards, the resin was pelleted at 6,000 x g and washed to remove non-specific bound proteins. FLAG-fusion proteins were eluted by incubating the resin with 100 µL of free FLAG peptide at a concentration of 0.5 mg/mL for 15 min, three times. The pooled elution was either provided to the Mass Spectrometry Core Facility at Johns Hopkins University or prepared for western blot analysis.

### Bacterial Two-Hybrid

Plasmids containing fusions to T18 and T25 were co-transformed into BTH101 competent cells and plated onto LB agar supplemented with ampicillin (100 μg/μL) and kanamycin (50 μg/μL) and grown overnight at 30°C. Biological triplicates of each co-transformation were grown overnight at 30°C in LB including with ampicillin (100 μg/μL), kanamycin (50 μg/μL), and IPTG (0.5 mM). After overnight incubation, 2 µL of each was spotted on LB agar plates supplemented with ampicillin (100 µg/µL), kanamycin (50 µg/µL), X-gal (40 μg/mL), and IPTG (0.5 mM) and incubated for 2 days at 30°C. Blue colonies were considered positive protein-protein interactions.

### DNA damage quantification using the *P_sidA_-egfp* reporter construct

To quantify DNA damage using the *P_sidA_-egfp* reporter, average fluorescence intensity for each cell in three biological replicate experiments per strain was measured using FIJI with the MicrobeJ plugin. We calculated “High GFP” intensity, which we considered above the 99^th^ percentile of the specified control strain GFP values, by first sorting the average fluorescent intensities for each replicate from lowest to highest. For the *fzlA* over-expression or deletion and *ftsW**I\** strains, we compared fluorescent intensity values to the 99^th^ percentile of WT GFP values. For the FtsK depletion strains we compared fluorescent intensity values to the 99^th^ percentile of the replete condition of the FtsK_FL_ depletion strain in a WT background grown for 4 hours in the presence of 0.3% xylose. We then multiplied the total number of cells analyzed for each control replicate by 0.99. The average intensity of that resulting cell number was then recorded. After obtaining the fluorescence intensity of the 99^th^ percentile of each control replicate, we averaged the fluorescence intensity of the three replicates. For all of the strains analyzed we looked for fluorescence intensity values above the calculated 99^th^ percentile of the control strain average. The number of cells that had higher values were counted and then divided by the total number of cells analyzed for each replicate of each strain. The average percentage of cells above the 99^th^ percentile of the control strain fluorescent intensity was then calculated by averaging the percentages of each replicate for each strain. These average percentages were then used for graphing and subsequent analysis.

## Acknowledgements

The authors would like to thank Lucy Shapiro, Michael Laub, Régis Hallez, and Patrick Viollier for strains and antibodies and Luke Lavis for Janelia Fluor dyes. We thank members of the Goley and Xiao laboratories for helpful discussions, especially Zhixin (Jason) Lyu and we thank Erika Smith for helpful comments on this manuscripts. This study was supported by the National Institutes of Health under awards R35GM136221 (EDG), R01GM108640 (EDG), R35GM136436 (JX), and T32GM144272 (training grant support of IPP) and by the National Natural Science Foundation of China (award 32270035 to XY) and Provincial Natural Science Foundation of Anhui (award 2208085MC40 to XY).

## Supplemental Materials Summary

Fig. S1 contains western blots that confirm the phenotypes and protein levels of various strains used throughout this article. Fig. S2 demonstrates example trajectories and speed measurements generated by SMT of Halo-FtsW. Fig. S3 illustrates that ZapA-mNeongreen is produced and localizes to the division plane, validating its use for locating Z-rings in FzlA depleted cells. Fig S4 is a reanalysis of the single molecule tracking from Fig. 3-4, similar to the re-analysis displayed in Fig. 5, to demonstrate that modulating FzlA levels can modulate the state of the two-populations of FtsW dynamics. Fig. S5 summarizes the tested combinations of FzlA and FtsK variants in each orientation for interaction by BTH. Table S1 summarizes the SMT results of Halo-FtsW and Halo-FzlA in the various backgrounds and conditions referred to throughout the text. Table S2 summarizes the results from mass spectrometry-based protein identification of eluent from the 3xFLAG-FzlA and wild-type co-immunoprecipitation samples. Table S3 summarizes the DNA-damage results from the various strains and conditions referred to in Fig. 9. Video 1 is an example timelapse of a *Caulobacter* cell constricting and dividing. Videos 2-4 are examples of single molecules of FtsW** (Video 2) and FtsW (Videos 3-4) that illustrate example movements by single-molecule tracking. Videos 5-7 are examples of the three constriction defects (Video 5, both daughters halt growth post-division. Video 6, one daughter halts growth post-division. Video 7, one daughter lyses post-division) observed during FzlA overproduction.

## Supplementary figure, table, and video legends

**Figure S1.**
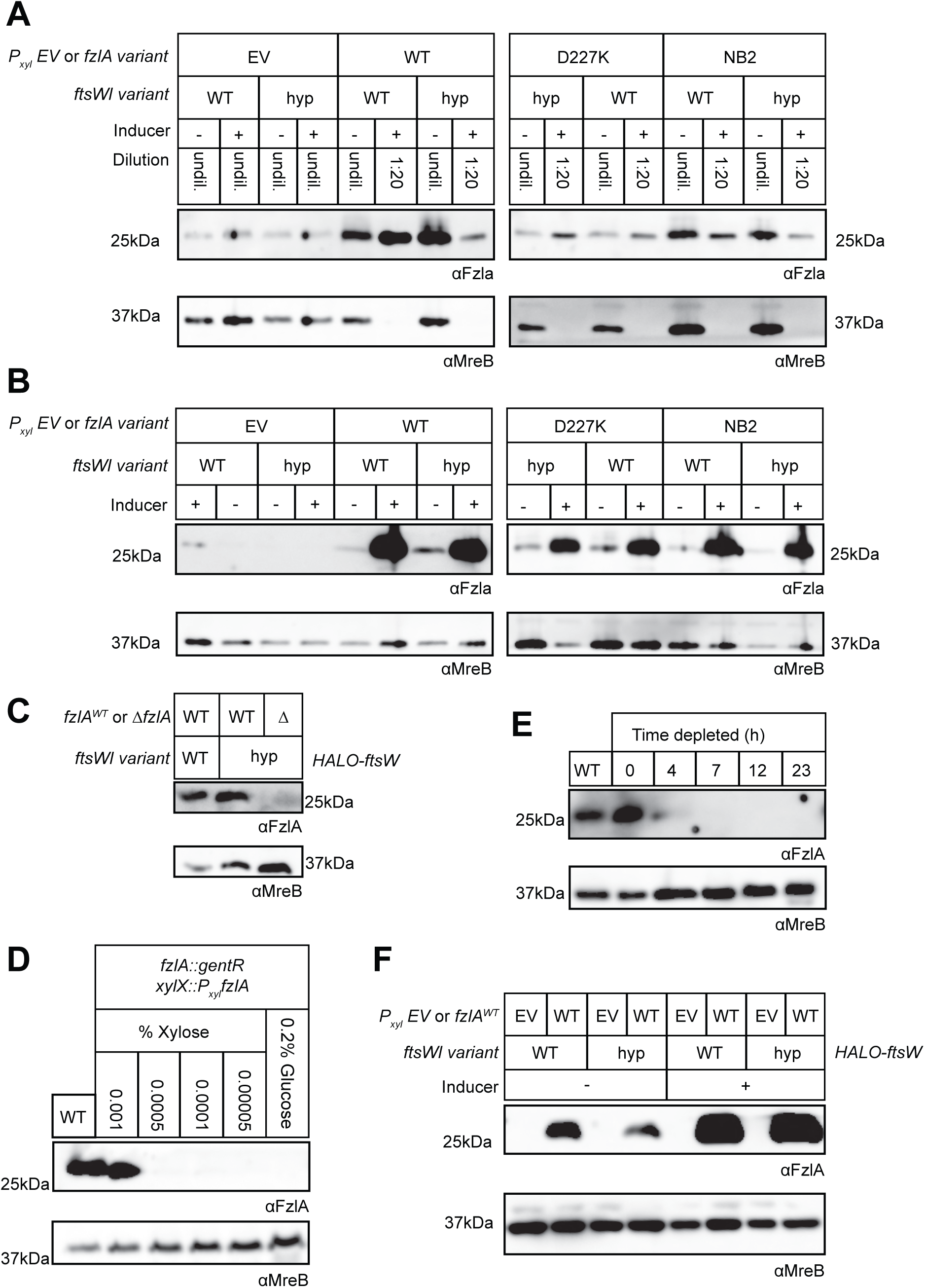
Western blots to confirm FzlA levels during strain characterization. **A-F.** Western blot analysis using primary antibody recognizing either FzlA (αFzlA) or the loading control MreB (αMreB). Ladder values next to blots are in kilodaltons (kDa). **A-B.** *fzlA* variant overexpression in both wild-type (WT) and *ftsW**I\** backgrounds with empty vector (EV) controls. **A.** Strains with high FzlA were diluted 1:20 before loading as indicated to enable detection of all samples at a single exposure time. **B.** Same as **A** but samples were all undiluted. **C.** FzlA is not detected in the Δ*fzlA; ftsW**I\** strain. **D.** *fzlA* induction in the *halo-ftsW* background (EG3523) as a function of xylose concentration to match FzlA levels in the wild-type *halo-ftsW* (EG3052) background. We selected 0.001% xylose as the closest approximation of FzlA production without underproducing. **E.** FzlA depletion in the *halo-ftsW* background (EG3523). **F.** FzlA over-production in the *halo-ftsW* (EG3519) and *halo-ftsW\*\**; *ftsI\** (EG3525) backgrounds, as well as the respective empty vector (EV) controls (WT: EG3537. *ftsW**I\**: EG3538).

**Figure S2.**
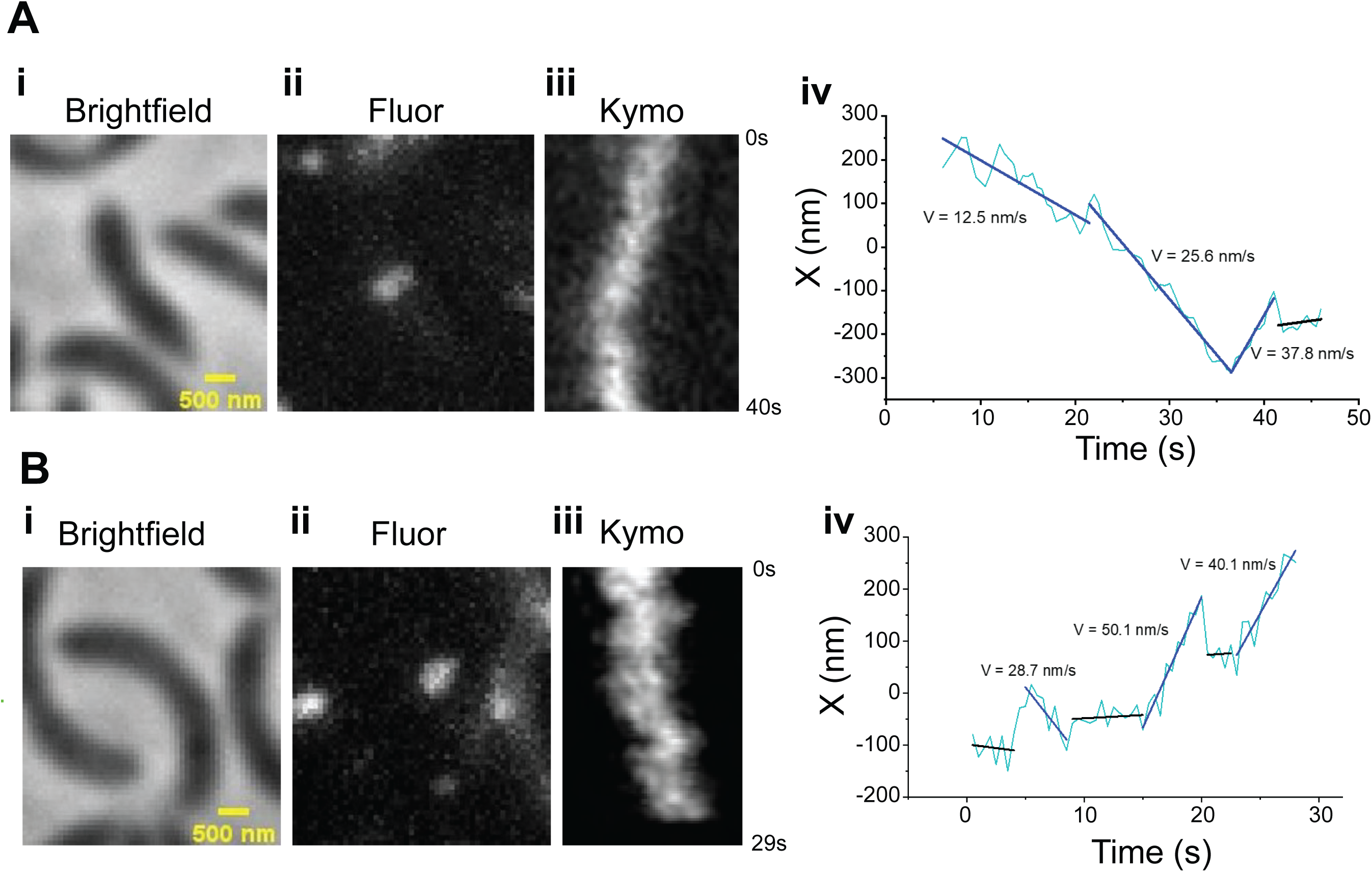
Example trajectories of single molecules of Halo-FtsW showing FtsW may change speeds. **A-B. A. i.** Brightfield image of a *halo-ftsW* background cell (EG3052). **ii.** Representative maximum fluorescence intensity projection image (Fluor) for single labeled Halo-FtsW. **iii.** Kymograph (Kymo) of the fluorescence signal of a line scan across the division plane that encapsulates a labeled single molecule of Halo-FtsW**. **iv.** Plot of the molecule position within the line scan over the course of single molecule tracking.

**Figure S3.**
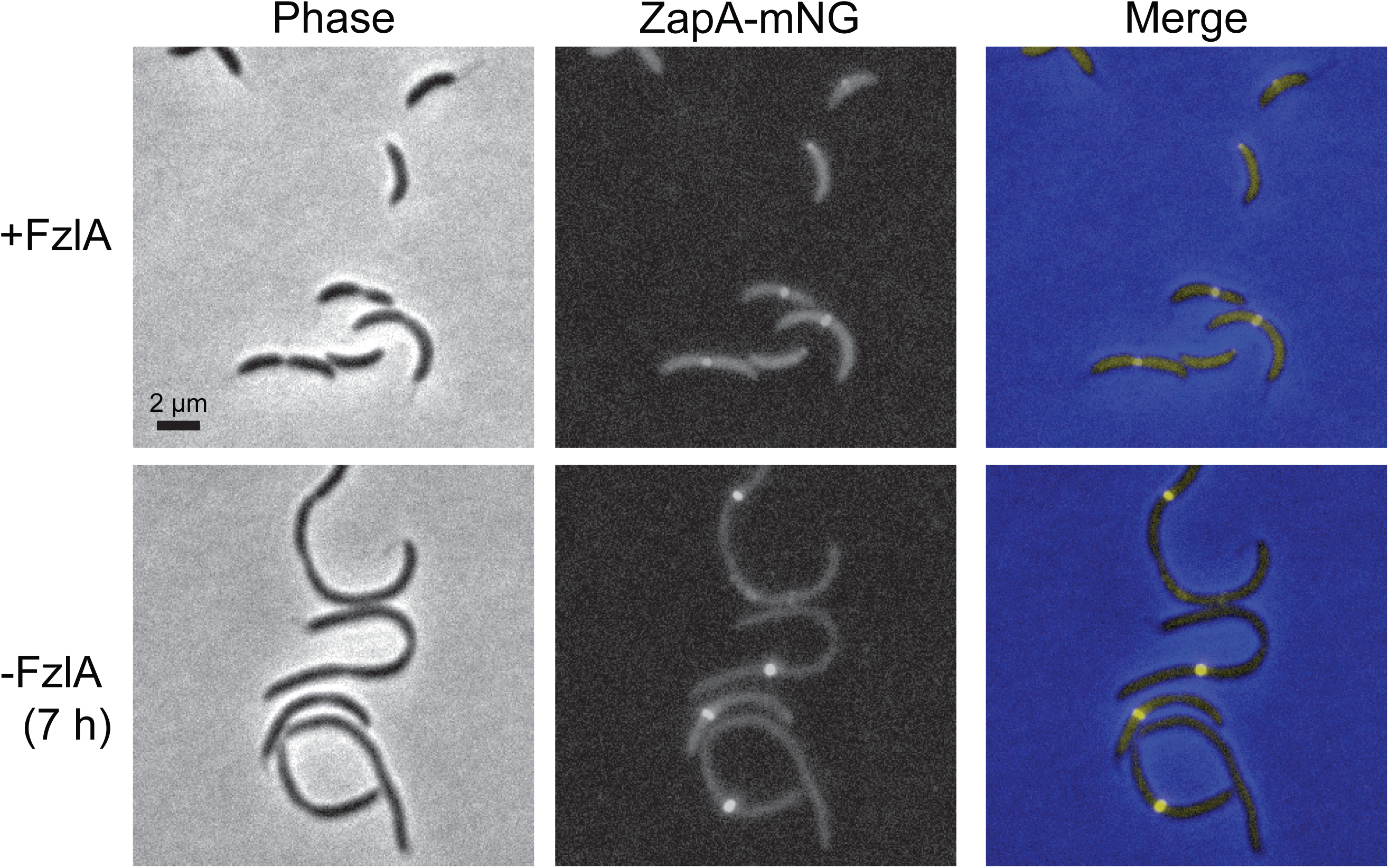
ZapA localization reflects Z-ring position in filamentous cells depleted of FzlA. Phase contrast, epifluorescence, and merged images of a strain expressing *zapA-mNG* dependent on xylose for production of FzlA (EG3523). The strain was either continuously induced (top) or depleted for seven hours (bottom) by being grown in the presence of 0.001% xylose or glucose, respectively.

**Figure S4.**
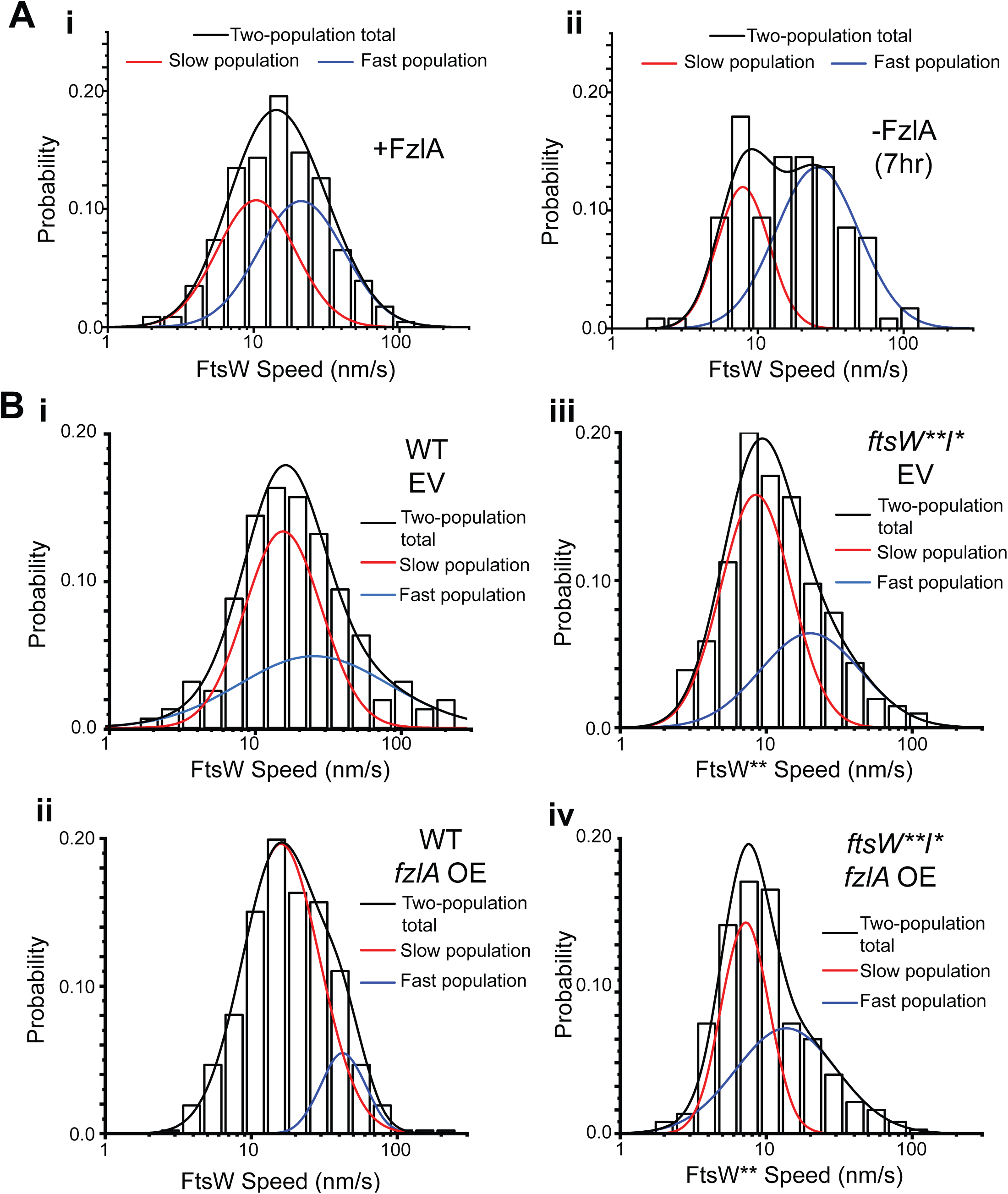
FzlA converts fast-moving FtsW into a slow-moving, active state. **A.** Histograms of SMT data for Halo-FtsW (WT or FtsW**) with calculated two-population curves that best fit the data for **i.** FtsW in cells producing FzlA (+FzlA, 0.001% xylose) or **ii.** depleted of FzlA for seven hours (-FzlA (7 h), 0.001% glucose) (re-analysis of data presented in Fig. 3A). **Bi-ii.** FtsW in **i.** a *halo-ftsW* (WT EV): EG3537), **ii.** *fzlA* OE (WT +FzlA: EG3519), (re-analysis of data presented in Fig. 4A) **Biii-iv.** FtsW** in **iii.** *halo-ftsW\*\**; *ftsI\** (*ftsW**I\** EV: EG3538), **iv.** *fzlA OE* (*ftsW**I\** +FzlA: EG3525) background (re-analysis of data presented in Fig. 4C). Red = slow population. Blue = fast population. Black: total based on two-population.

**Figure S5.**
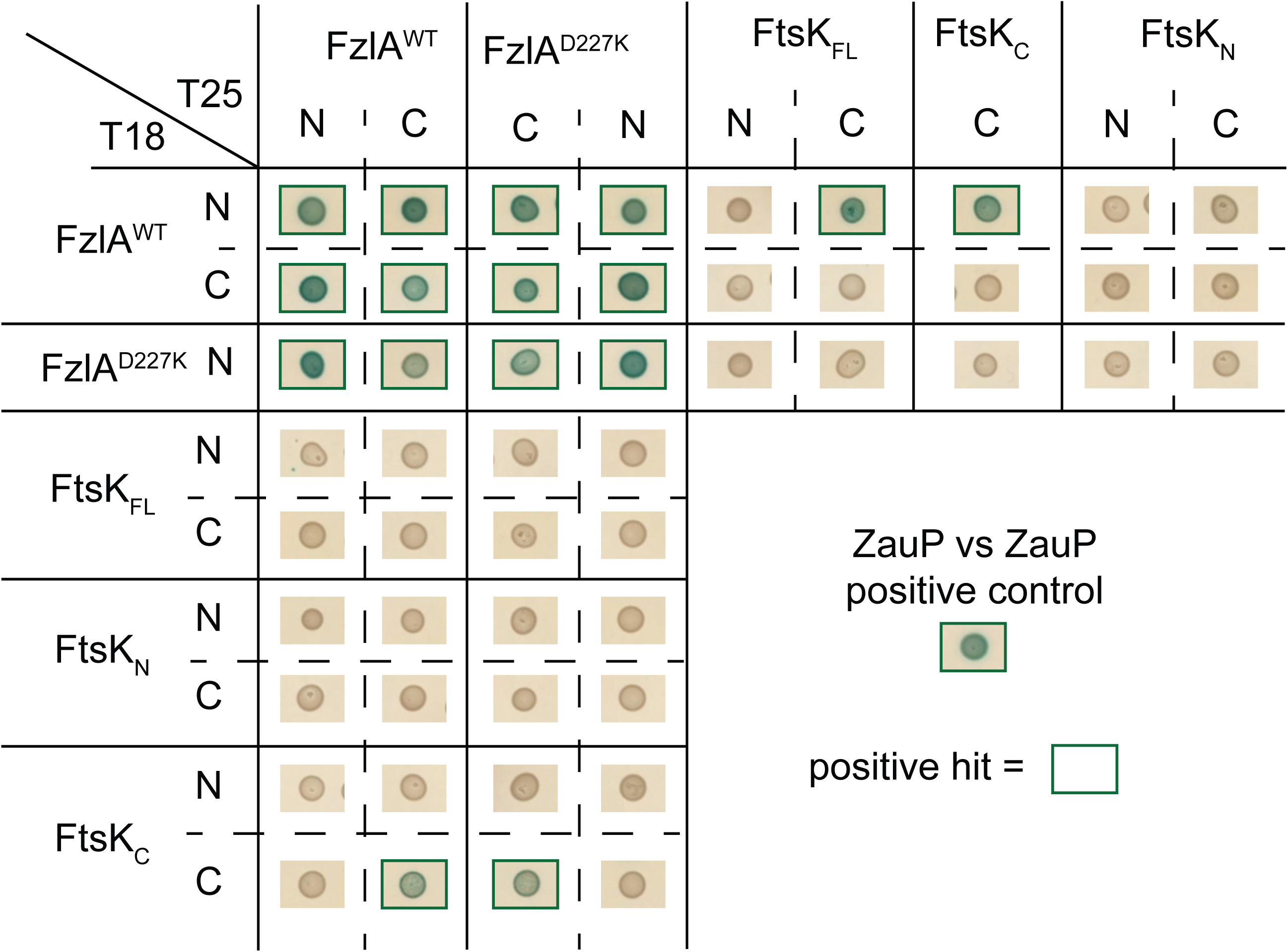
Complete results of bacterial two-hybrid analysis. Bacterial Two-Hybrid results for interaction between FzlA variants and full-length FtsK or its domains. The adenylyl cyclase subunits T18 (left) and T25 (top) are fused to proteins at the indicated terminus for each row or column. A green box around the representative spot image means that the three biological triplicates were positive for induction of the cAMP-dependent β-galactosidase reporter, indicating a positive interaction. FtsK_FL_, Full-length FtsK. FtsK_N_, FtsK N-domain. FtsK_C_, FtsK C-domain.

**Table S1. Summary of SMT of Halo-FtsW and Halo-FzlA variants with varying states of activation.** Average FtsW or FzlA molecule speeds (nm/s) are indicated with standard error of the mean (SEM), proportion of moving molecules (%), and the life-time of stationary FtsW (s). N: sample size. WT: wild-type. OE: overexpressed. EV: empty vector control. ++FzlA: ∼20-fold overproduced FzlA. +FzlA: induced to wild-type levels. -FzlA: without induction.

**Table S2. Proteins identified by mass-spectrometry that are five-fold enriched in relative abundance in 3xFLAG-FzlA eluate compared to WT.** Proteins identified by Mass-spectrometry with accession numbers, and information about relative abundance between 3xFLAG-Fzla and WT eluate. WT: wild-type. kDa: kilodalton. --: not detected in WT sample.

**Table S3. Summary of DNA damage reporter experiments.** Percentage (%) of cells with high levels of *P*_sidA_-driven EGFP production in backgrounds with varying FzlA or FtsK levels and states of FtsWI activation. WT: wild-type. NA: Not applicable (not an inducible strain). OE: overexpression. EV: empty vector control. ++FzlA: ∼20-fold overproduced FzlA. FtsK_C_: FtsK C-terminus. FtsK_FL_: FtsK full-length.

**Table S4. Strains and plasmids used in this study.**

**Video 1. A wild-type division that completes successfully.** Representative timelapse of a constricting wild-type cell with an empty vector control. Each frame is 5 minutes apart and the total timelapse is 4 hours.

**Video 2. FtsW** movement about the division plane in a *ftsW**I\** background.** Representative timelapse of fluorescent JF646-labeled Halo-FtsW** in a *ftsW**I\** background over 140 s. This timelapse corresponds to the kymograph present in Figures 2aiii. Each frame is 500 ms.

**Video 3. FtsW moving at multiple speeds in a wild-type background.** Representative timelapse of fluorescent JF646-labeled Halo-FtsW in a wild-type background over 40 s. This timelapse corresponds to the kymograph present in Figure S2aiii. Each frame is 500 ms.

**Video 4. FtsW moving at multiple speeds in a wild-type background.** Representative timelapse of fluorescent JF646-labeled Halo-FtsW in a wild-type background over 29 s. This timelapse corresponds to the kymograph present in Figure S2aiii. Each frame is 500 ms.

**Video 5. A division in which both daughter cells halt growth post-constriction.** Representative timelapse of a FzlA overproducing cell in a WT background for which both daughter cells halt growth post-division. Each frame is 5 minutes apart and the total timelapse is 4 hours.

**Video 6. A division in which one daughter cell halts growth post-constriction.** Representative timelapse of a FzlA-overproducing cell in a WT background for which one daughter cell halts growth post-division. Each frame is 5 minutes apart and the total timelapse is 4 hours.

**Video 7. A division in which one daughter cell lyses growth post-constriction.** Representative timelapse of a FzlA overproducing cell in a WT background for which one daughter cell lyses post-division. Each frame is 5 minutes apart and the total timelapse is 4 hours.

## References

Barrows, J.M., and E.D. Goley. 2021. FtsZ dynamics in bacterial division: What, how, and why? Curr Opin Cell Biol. 68:163–172. doi:10.1016/j.ceb.2020.10.013.

Beaufay, F., Coppine, J., Mayard, A., Laloux, G., De Bolle, X., and Hallez, R. 2015. A NAD-dependent glutamate dehydrogenase coordinates metabolism with cell division in *Caulobacter crescentus*. The EMBO J. 34: 1786–1800. doi:10.15252/embj.201490730.

Bernhardt, T.G., and P.A.J. de Boer. 2004. Screening for synthetic lethal mutants in Escherichia coli and identification of EnvC (YibP) as a periplasmic septal ring factor with murein hydrolase activity. Mol Microbiol. 52:1255–1269. doi:10.1111/j.1365-2958.2004.04063.x.

Bernhardt, T.G., and P.A.J. de Boer. 2005. SlmA, a nucleoid-associated, FtsZ binding protein required for blocking septal ring assembly over chromosomes in E. coli. Mol Cell. 18:555–564. doi:10.1016/j.molcel.2005.04.012.

Bigot, S., V. Sivanathan, C. Possoz, F.X. Barre, and F. Cornet. 2007. FtsK, a literate chromosome segregation machine. Mol Microbiol. 64:1434–1441. doi:10.1111/j.1365-2958.2007.05755.x.

Bisson-Filho, A.W., Y.-P. Hsu, G.R. Squyres, E. Kuru, F. Wu, C. Jukes, Y. Sun, C. Dekker, S. Holden, M.S. Vannieuwenhze, Y. v Brun, and E.C. Garner. 2017. Treadmilling by FtsZ filaments drives peptidoglycan synthesis and bacterial cell division. Science. 355:739–743. doi:10.1126/science.aak9973.

Chen, J.C., Viollier, P.H., and Shapiro, L. 2005. A membrane metalloprotease participates in the sequential degradation of a *Caulobacter* polarity determinant. Mol Microbiol. 55: 1085–1103. doi:10.1111/j.1365-2958.2004.04443

Coltharp, C., and J. Xiao. 2017. Beyond force generation: Why is a dynamic ring of FtsZ polymers essential for bacterial cytokinesis? Bioessays. 39:1–11. doi:10.1111/mec.13536.Application.

Daitch, A.K., and E.D. Goley. 2020. Uncovering Unappreciated Activities and Niche Functions of Bacterial Cell Wall Enzymes. Current Biology. 30:R1170–R1175. doi:10.1016/j.cub.2020.07.004.

Dewachter, L., N. Verstraeten, M. Fauvart, and J. Michiels. 2018. An integrative view of cell cycle control in Escherichia coli. FEMS Microbiol Rev. 42:116–136. doi:10.1093/femsre/fuy005.

Draper, G.C., N. Mclennan, K. Begg, M. Masters, and W.D. Donachie. 1998. Only the N-Terminal Domain of FtsK Functions in Cell Division. J Bacteriol. 180:4621–4627. Doi:10.1128/JB.180.17.4621-4627.1998.

Du, S., S. Pichoff, and J. Lutkenhaus. 2016. FtsEX acts on FtsA to regulate divisome assembly and activity. Proceedings of the National Academy of Sciences. 113:E5052–E5061. doi:10.1073/pnas.1606656113.

Gerding, M.A., B. Liu, F.O. Bendezú, C.A. Hale, T.G. Bernhardt, and P.A.J. de Boer. 2009. Self-enhanced accumulation of FtsN at division sites and roles for other proteins with a SPOR domain (DamX, DedD, and RlpA) in Escherichia coli cell constriction. J Bacteriol. 191:7383–7401. doi:10.1128/JB.00811-09.

Goley, E.D., Dye, N.A., Werner, J.N., Gitai, Z., and L. Shapiro. 2010. Imaging-based identification of a critical regulator of FtsZ protofilament curvature in Caulobacter. Mol Cell. 144:724–732. doi:10.1038/jid.2014.371.

Goley, E.D., Y. Yi-Chun, S.-H. Hong, M.J. Fero, E. Abeliuk, H.H. McAdams, and L. Shapiro. 2011. Assembly of the Caulobacter cell division machine. Mol Microbiol. 9:19–22. doi:10.3816/CLM.2009.n.003.Novel.

Grimm, J.B., A.K. Muthusamy, Y. Liang, T.A. Brown, W.C. Lemon, R. Patel, R. Lu, J.J. Macklin, P.J. Keller, N. Ji, and L.D. Lavis. 2017. A general method to fine-tune fluorophores for live-cell and in vivo imaging. Nature Methods 2017 14:10. 14:987–994. doi:10.1038/nmeth.4403.

Heidrich, C., M.F. Templin, A. Ursinus, M. Merdanovic, J. Rgen Berger, H. Schwarz, M.A. de Pedro, and J.-V. Hö. 2001. Involvement of N-acetylmuramyl-L-alanine amidases in cell separation and antibiotic-induced autolysis of Escherichia coli. Mol Micro. 41:167–178. Doi:10.1046/j.1365-2958.2001.02499.x.

Lambert, A., A. Vanhecke, A. Archetti, S. Holden, F. Schaber, Z. Pincus, M.T. Laub, E. Goley, and S. Manley. 2018. Constriction Rate Modulation Can Drive Cell Size Control and Homeostasis in C. crescentus. iScience. 4:180–189. doi:10.1016/j.isci.2018.05.020.

Lariviere, P.J., C.R. Mahone, G. Santiago-Collazo, M. Howell, A.K. Daitch, R. Zeinert, P. Chien, P.J.B. Brown, and E.D. Goley. 2019. An Essential Regulator of Bacterial Division Links FtsZ to Cell Wall Synthase Activation. Current Biology. 29:1460–1470.e4. doi:10.1016/j.cub.2019.03.066.

Lariviere, P.J., P. Szwedziak, C.R. Mahone, J. Löwe, and E.D. Goley. 2018. FzlA, an essential regulator of FtsZ filament curvature, controls constriction rate during Caulobacter division. Mol Microbiol. 107:180–197. doi:10.1111/mmi.13876.

Li, Y., A. Boes, Y. Cui, S. Zhao, Q. Liao, H. Gong, E. Breukink, J. Lutkenhaus, M. Terrak, and S. Du. 2021. Identification of the potential active site of the septal peptidoglycan polymerase FtsW. PLoS Genet. 18. doi:10.1371/journal.pgen.1009993.

Liu, B., L. Persons, L. Lee, and P.A.J. de Boer. 2015. Roles for both FtsA and the FtsBLQ subcomplex in FtsN-stimulated cell constriction in Escherichia coli. Mol Microbiol. 95:945– 970. doi:10.1111/mmi.12906.

Lord, S.J., K.B. Velle, R. Dyche Mullins, and L.K. Fritz-Laylin. 2020. SuperPlots: Communicating reproducibility and variability in cell biology. Journal of Cell Biology. 219. doi:10.1083/JCB.202001064.

Lyu, Z., A. Yahashiri, X. Yang, J.W. McCausland, G.M. Kaus, R. McQuillen, D.S. Weiss, and J. Xiao. 2022. FtsN maintains active septal cell wall synthesis by forming a processive complex with the septum-specific peptidoglycan synthases in E. coli. Nat Commun. 13:5751. doi:10.1038/s41467-022-33404-8.

Mahone, C.R., and E.D. Goley. 2020. Bacterial cell division at a glance. J Cell Sci. 133: jcs237057. doi:10.1242/jcs.237057.

Marmont, L.S., and T.G. Bernhardt. 2020. A conserved subcomplex within the bacterial cytokinetic ring activates cell wall synthesis by the FtsW-FtsI synthase. Proc Natl Acad Sci U S A. 117:23879–23885. doi:10.1073/pnas.2004598117.

McCausland, J.W., X. Yang, G.R. Squyres, Z. Lyu, K.E. Bruce, M.M. Lamanna, B. Söderström, E.C. Garner, M.E. Winkler, J. Xiao, and J. Liu. 2021. Treadmilling FtsZ polymers drive the directional movement of sPG-synthesis enzymes via a Brownian ratchet mechanism. Nat Commun. 12. doi:10.1038/s41467-020-20873-y.

McQuillen, R., and J. Xiao. 2020. Insights into the Structure, Function, and Dynamics of the Bacterial Cytokinetic FtsZ-Ring. Annu Rev Biophys. 49:309–341. doi:10.1146/annurev-biophys-121219-081703.

Meier, E.L., A.K. Daitch, Q. Yao, A. Bhargava, G.J. Jensen, and E.D. Goley. 2017. FtsEX-mediated regulation of the final stages of cell division reveals morphogenetic plasticity in Caulobacter crescentus. PLoS Genetics. 13:e1006999. Doi:10.1371/journal.pgen.1006999.

Modell, J.W., A.C. Hopkins, and M.T. Laub. 2011. A DNA damage checkpoint in Caulobacter crescentus inhibits cell division through a direct interaction with FtsW. Genes Dev. 25:1328–1343. doi:10.1101/gad.2038911.a.

Modell, J.W., T.K. Kambara, B.S. Perchuk, and M.T. Laub. 2014. A DNA Damage-Induced, SOS-Independent Checkpoint Regulates Cell Division in Caulobacter crescentus. PLoS Biol. 12:e1001977. doi:10.1371/journal.pbio.1001977.

Park, K.T., S. Du, and J. Lutkenhaus. 2020. Essential role for ftsL in activation of septal peptidoglycan synthesis. mBio. 11:e03012–20. doi:10.1128/mBio.03012-20.

Park, K.T., S. Pichoff, S. Du, and J. Lutkenhaus. 2021. FtsA acts through FtsW to promote cell wall synthesis during cell division in Escherichia coli. Proc Natl Acad Sci U S A. 118:e2107210118. doi:10.1073/PNAS.2107210118/-/DCSUPPLEMENTAL.

Perez, A.J., Y. Cesbron, S.L. Shaw, J. Bazan Villicana, H.-C.T. Tsui, M.J. Boersma, Z.A. Ye, Y. Tovpeko, C. Dekker, S. Holden, and M.E. Winkler. 2019. Movement dynamics of divisome proteins and PBP2x:FtsW in cells of Streptococcus pneumoniae. Proceedings of the National Academy of Sciences. 116:3211–3220. doi:10.1073/pnas.1816018116.

Peters, N.T., T. Dinh, and T.G. Bernhardt. 2011. A Fail-safe mechanism in the septal ring assembly pathway generated by the sequential recruitment of cell separation amidases and their activators. J Bacteriol. 193:4973–4983. doi:10.1128/JB.00316-11.

lo Sciuto, A., R. Fernández-Piñar, L. Bertuccini, F. Iosi, F. Superti, and F. Imperi. 2014. The periplasmic protein TolB as a potential drug target in Pseudomonas aeruginosa. PLoS One. 9:e103784. doi:10.1371/journal.pone.0103784.

Squyres, G.R., M.J. Holmes, S.R. Barger, B.R. Pennycook, J. Ryan, V.T. Yan, and E.C. Garner. 2021. Single-molecule imaging reveals that Z-ring condensation is essential for cell division in Bacillus subtilis. Nat Microbiol. 6:553–562. doi:10.1038/s41564-021-00878-z.

Tsang, M.J., and T.G. Bernhardt. 2015. A role for the FtsQLB complex in cytokinetic ring activation revealed by an ftsL allele that accelerates division. Mol Microbiol. 95:925–944. doi:10.1111/mmi.12905.

Uehara, T., K.R. Parzych, T. Dinh, and T.G. Bernhardt. 2010. Daughter cell separation is controlled by cytokinetic ring-activated cell wall hydrolysis. EMBO Journal. 29:1412–1422. doi:10.1038/emboj.2010.36.

Wang, L., and J. Lutkenhaus. 1998. FtsK is an essential cell division protein that is localized to the septum and induced as part of the SOS response. Mol Microbiol. 29:731–740. doi:10.1046/j.1365-2958.1998.00958.x.

Wang, S.C.E., L. West, and L. Shapiro. 2006. The bifunctional FtsK protein mediates chromosome partitioning and cell division in Caulobacter. J Bacteriol. 188:1497–1508. doi:10.1128/JB.188.4.1497-1508.2006.

Weiss, D.S. 2015. Last but not least: New insights into how FtsN triggers constriction during Escherichiacoli cell division. Mol Microbiol. 95:903–909. doi:10.1111/mmi.12925.

Woldemeskel, S.A., R. McQuillen, A.M. Hessel, J. Xiao, and E.D. Goley. 2017. A conserved coiled-coil protein pair focuses the cytokinetic Z-ring in Caulobacter crescentus. Mol Microbiol. 105:721–740. doi:10.1111/mmi.13731.

Yahashiri, A., M.A. Jorgenson, and D.S. Weiss. 2015. Bacterial SPOR domains are recruited to septal peptidoglycan by binding to glycan strands that lack stem peptides. Proceedings of the National Academy of Sciences. 112:11347–11352. doi:10.1073/pnas.1508536112.

Yang, X., and R. Liu. 2022. How does FtsZ’s treadmilling help bacterial cells divide? BIOCELL. 46:2343–2351. doi:10.32604/BIOCELL.2022.022100.

Yang, X., Z. Lyu, A. Miguel, R. McQuillen, K.C. Huang, and J. Xiao. 2017. GTPase activity– coupled treadmilling of the bacterial tubulin FtsZ organizes septal cell wall synthesis. Science (1979). 355:744–747. doi:10.1126/science.aak9995.

Yang, X., R. McQuillen, Z. Lyu, P. Phillips-Mason, A. de La Cruz, J.W. McCausland, H. Liang, K.E. DeMeester, C.C. Santiago, C.L. Grimes, P. de Boer, and J. Xiao. 2021. A two-track model for the spatiotemporal coordination of bacterial septal cell wall synthesis revealed by single-molecule imaging of FtsW. Nat Microbiol. 6:584–593. doi:10.1038/s41564-020-00853-0.

Yu, X.-C., E.K. Weihe, and W. Margolin. 1998. Role of the C Terminus of FtsK in Escherichia coli Chromosome Segregation. 180. 6424–6428 pp.

## Supplemental Material References

Barrows, J. M., Sundararajan, K., Bhargava, A., & Goley, E. D. (2020). FtsA regulates z-ring morphology and cell wall metabolism in an FtsZ C-terminal linker-dependent manner in caulobacter crescentus. Journal of Bacteriology, 202(7). https://doi.org/10.1128/JB.00693-19

Battesti, A., & Bouveret, E. (2012). The bacterial two-hybrid system based on adenylate cyclase reconstitution in Escherichia coli. Methods, 58(4), 325–334. https://doi.org/10.1016/J.YMETH.2012.07.018

Evinger, M., & Agabian, N. (1977). Envelope-Associated Nucleoid from Caulobacter crescentus Stalked and Swarmer Cells. In JOURNAL OF BACTERIOLOGY. https://journals.asm.org/journal/jb

Goley, E. D. (2010). Imaging-based identification of a critical regulator of FtsZ protofilament curvature in Caulobacter. Mol Cell, 144(5), 724–732. https://doi.org/10.1038/jid.2014.371

Goley, E. D., Yi-Chun, Y., Hong, S.-H., Fero, M. J., Abeliuk, E., McAdams, H. H., & Shapiro, L. (2011). Assembly of the Caulobacter cell division machine. Molecular Microbiology, 9(1), 19–22. https://doi.org/10.3816/CLM.2009.n.003.Novel

Lariviere, P. J., Mahone, C. R., Santiago-Collazo, G., Howell, M., Daitch, A. K., Zeinert, R., Chien, P., Brown, P. J. B., & Goley, E. D. (2019). An Essential Regulator of Bacterial Division Links FtsZ to Cell Wall Synthase Activation. Current Biology, 29(9), 1460–1470.e4. https://doi.org/10.1016/j.cub.2019.03.066

Lariviere, P. J., Szwedziak, P., Mahone, C. R., Löwe, J., & Goley, E. D. (2018). FzlA, an essential regulator of FtsZ filament curvature, controls constriction rate during Caulobacter division. Molecular Microbiology, 107(2), 180–197. https://doi.org/10.1111/mmi.13876

Modell, J. W., Hopkins, A. C., & Laub, M. T. (2011). A DNA damage checkpoint in Caulobacter crescentus inhibits cell division through a direct interaction with FtsW. Genes & Development, 25(12), 1328. https://doi.org/10.1101/gad.2038911.a

Modell, J. W., Kambara, T. K., Perchuk, B. S., & Laub, M. T. (2014). A DNA Damage-Induced, SOS-Independent Checkpoint Regulates Cell Division in Caulobacter crescentus. PLoS Biology, 12(10). https://doi.org/10.1371/journal.pbio.1001977

Thanbichler, M., Iniesta, A. A., & Shapiro, L. (2007). A comprehensive set of plasmids for vanillate - And xylose-inducible gene expression in Caulobacter crescentus. Nucleic Acids Research, 35(20). https://doi.org/10.1093/nar/gkm818

Wang, S. C. E., West, L., & Shapiro, L. (2006). The bifunctional FtsK protein mediates chromosome partitioning and cell division in Caulobacter. Journal of Bacteriology, 188(4), 1497–1508. https://doi.org/10.1128/JB.188.4.1497-1508.2006

